# Conkazal-M1 from the MKAVA family of conotoxins – a dual-function protease inhibitor and neuroactive peptide

**DOI:** 10.1101/2025.11.05.686841

**Authors:** Celeste M. Hackney, Thomas Lund Koch, Nicklas Lund Ryding, Aymeric Rogalski, Kevin Chase, Matías Leonel Giglio, Samuel S. Espino, Zildjian G. Acyatan, Maren Watkins, Baldomero M. Olivera, Helena Safavi-Hemami, Kaare Teilum, Lars Ellgaard

**Author notes:** **Correspondence:** Lars Ellgaard, University of Copenhagen, Department of Biology, Ole Maaløes Vej 5, DK-2200 Copenhagen N., Denmark., Kaare Teilum, University of Copenhagen, Department of Biology, Ole Maaløes Vej 5, DK-2200 Copenhagen N., Denmark.

## Abstract

Marine cone snails produce a diverse array of bioactive peptides, known as conotoxins, in their venom. Given their high target potency and specificity, conotoxins are attractive compounds for the development of precision research tools and pharmacological agents. Here, we provide the first experimental characterization of a conotoxin from the MKAVA superfamily, conkazal-M1, from *Conus magus*. Using NMR spectroscopy, we show that conkazal-M1 adopts a fold characteristic of the Kazal-type protease inhibitor family, featuring a Glu residue at the inhibitory P1 position. Recombinantly expressed conkazal-M1 inhibits the proteolytic activity of Subtilisin A with an apparent Ki of 1.1 μM. In addition, conkazal-M1 partially inhibits calcium transients in mouse sensory neurons, suggesting a potential role in modulating ion-channel activity, as seen for many other toxins. The dual function of conkazal-M1 in protease inhibition and neuroactivity is analogous to the dual function of several toxins harboring a Kunitz-type fold. The well-conserved sequence of the MKAVAs indicates an evolutionary trajectory in which these proteins face an adaptive conflict, where mutations that enhance one activity compromise the other. Collectively, this work provides new structural and functional insights into a previously uncharacterized toxin superfamily in cone snails, illustrates how structural scaffolds can be repurposed for functions that diverge from the original while retaining their overall structure, and expands our understanding of the toxin arsenal available to venomous animals.

## INTRODUCTION

Peptide toxins from animal venoms target a broad range of biological processes to subdue prey or deter predators (1). To exert these effects, toxins act by disrupting the cellular ion balance or by interfering with signaling pathways mediated by transmembrane proteins such as ion channels, G protein–coupled receptors (GPCRs), transporters, and tyrosine kinases (2). Other major classes of toxins target hemostatic processes, such as blood coagulation (3). Due to the evolutionary conservation of such key processes and pathways, many peptide toxins hold great potential as research tools and potential pharmaceutical compounds.

The diverse functions of peptide toxins are enabled by a variety of molecular structures. Many of these toxins fold into small, compact structures stabilized by multiple disulfide bonds. While the cysteine residues involved in disulfide-bond formation are typically conserved, inter-cysteine residues normally undergo rapid evolution that generates diversity (4, 5). Consequently, toxin sequences – even within the same evolutionary family – often exhibit a high degree of sequence variation.

Several prevalent toxins function as protease inhibitors, reflecting the critical role of protease activity in the physiology of target animals. For example, venom protease inhibitors target proteases in the blood-coagulation system (causing internal bleeding and hypotension), enzymes of the cholinergic system (disrupting neurotransmitter metabolism), and the immune system (modulating immune and inflammatory responses). Common protease-inhibitor folds present in venoms include Kazal-, Kunitz-, and cystatin-like domains, Snake Venom Serine Protease Inhibitors, and Cysteine-Rich Secretory Proteins. As is common for other venom peptides, such folds are often recruited into the venom gland from endogenous genes and are subsequently “weaponized” (6). Through this process, these compact and stable structural scaffolds can be repurposed for functions that diverge from the original, while retaining their overall structure.

Well-characterized Kunitz-type toxins are known to have evolved new functions that differ from their original role as protease-inhibitors. For instance, the conkunitzins from venomous marine cone snails (7, 8), as well as mambaquaretin-1 (9) and calcicludine (10) from the Eastern green mamba, have evolved to inhibit ion channels or receptors. Moreover, certain Kunitz-like toxins display a dual function as both protease inhibitors and ion channel blockers (11–14). At the molecular level, the dual function of the tarantula Kunitz-type toxin HWTX-X is achieved through two distinct binding interfaces (12). It has also been shown that Kunitz-type protease inhibitors from venomous snakes undergo positive selection at sites that cover 30-50% of the domain (15). These examples provide textbook illustrations of the sequence plasticity often observed in toxins that allows them to adopt new functions using a preexisting structural scaffold.

A prominent example of a protease-inhibitor fold found in several venoms is the Kazal-type domain that inhibits a broad range of serine proteases. Kazal-type venom peptides are found in various venomous animals, including assassin bugs, shrew, honeybees, and wasps, where they have been associated with functions like anti-coagulation (16–18), pro-coagulation (19), antimicrobial activity (18), melanization in insects (20), and analgesia (19). In humans, the SPINK family of Kazal-type protease inhibitors (KPIs) plays important functions in regulating cellular protease activity. This is exemplified by the pancreatic trypsin inhibitor SPINK1, in which sequence variants are associated with pancreatitis and pancreatic cancer (21). Another example of a well-characterized KPI is greglin from the locust *Schistocerca gregaria*, which inhibits both elastase and subtilisin with low nanomolar Ki-values (22). In general, KPIs function as substrate analogs. They act by inserting their inhibitory P1 residue, the residue positioned immediately adjacent to the scissile bond, into the specificity pocket of the target protease (23). This residue is typically located two positions C-terminally to the second cysteine in the sequence on the so-called reactive-site loop (23). The nature of the P1 residue can vary considerably, in turn influencing substrate specificity – for instance, trypsin-inhibiting KPIs often contain a lysine or arginine residue at the P1 position to mimic physiological trypsin substrates (23). Classical KPIs harbor 40-60 amino acid residues and fold into a structure that comprises an α-helix and a three-stranded β-sheet held together by three disulfide bonds (23). KPIs that deviate from this definition (e.g. those with more or fewer disulfides or varying disulfide patterns) are classified as non-classical KPIs (see Results for more details).

The conotoxins produced by venomous marine cone snails are peptides encoded as precursor sequences that comprise a well-conserved N-terminal signal sequence that directs the peptide to the endoplasmic reticulum, followed in some cases by a propeptide of intermediate sequence conservation, and, finally, the mature conotoxin sequence. Approximately 70 different conotoxin gene families have been identified to date (24–26). While most mature conotoxins are 15-45 residues in length, many larger peptides also exist (25). Many families of large conotoxins – also known as macro-conotoxins (27) – remain uncharacterized due to the difficulty in producing large disulfide-bonded peptides. These larger peptides are expected to perform functions unavailable to shorter conotoxins, e.g., acting as enzymes or interacting with their targets through a larger binding interface.

Here, we surveyed venom gland transcriptomes to identify new macro-conotoxin families. We describe the previously uncharacterized MKAVA family of conotoxins and show by NMR spectroscopy that an MKAVA-family toxin from *Conus magus*, conkazal-M1, comprises a single non-classical Kazal-type domain with both protease inhibitor activity and the ability to inhibit KCl-induced calcium influx in a subset of dorsal root ganglia (DRG). We also uncover the evolutionary origin of the MKAVA-family toxins and discuss potential mechanisms that shaped their molecular evolution.

## RESULTS

### The MKAVA conotoxin superfamily is widely distributed among venomous cone snails

To characterize the MKAVA conotoxin superfamily, we mined publicly available venom gland transcriptomes from 49 cone snail species and identified 89 transcripts sharing the MKAVA consensus signal sequence (MKAVAVFLVVALAVAYG) with slight variations (Figure S1; all sequences provided in Supplementary File 1). As illustrated in Figure 1 and S1, sequences from snails with a prey preference from all three feeding groups of venomous snails (fish, other molluscs, and worms) are present among the analyzed sequences. In addition to their wide distribution across *Conus* species, we found that closely related MKAVA-like peptides are also found in other members of Conoidea, the larger superfamily of marine snails to which cone snails belong. Searches against the NCBI Sequence Read Archive (SRA) database identified similar sequences in species belonging to Drillidae, Turridae, Profundiconus, and Terebridae (see Materials and Methods section for SRA entry numbers). It should be noted that these sequences are incomplete, as they derive from unassembled short-read datasets.

**Figure 1.**
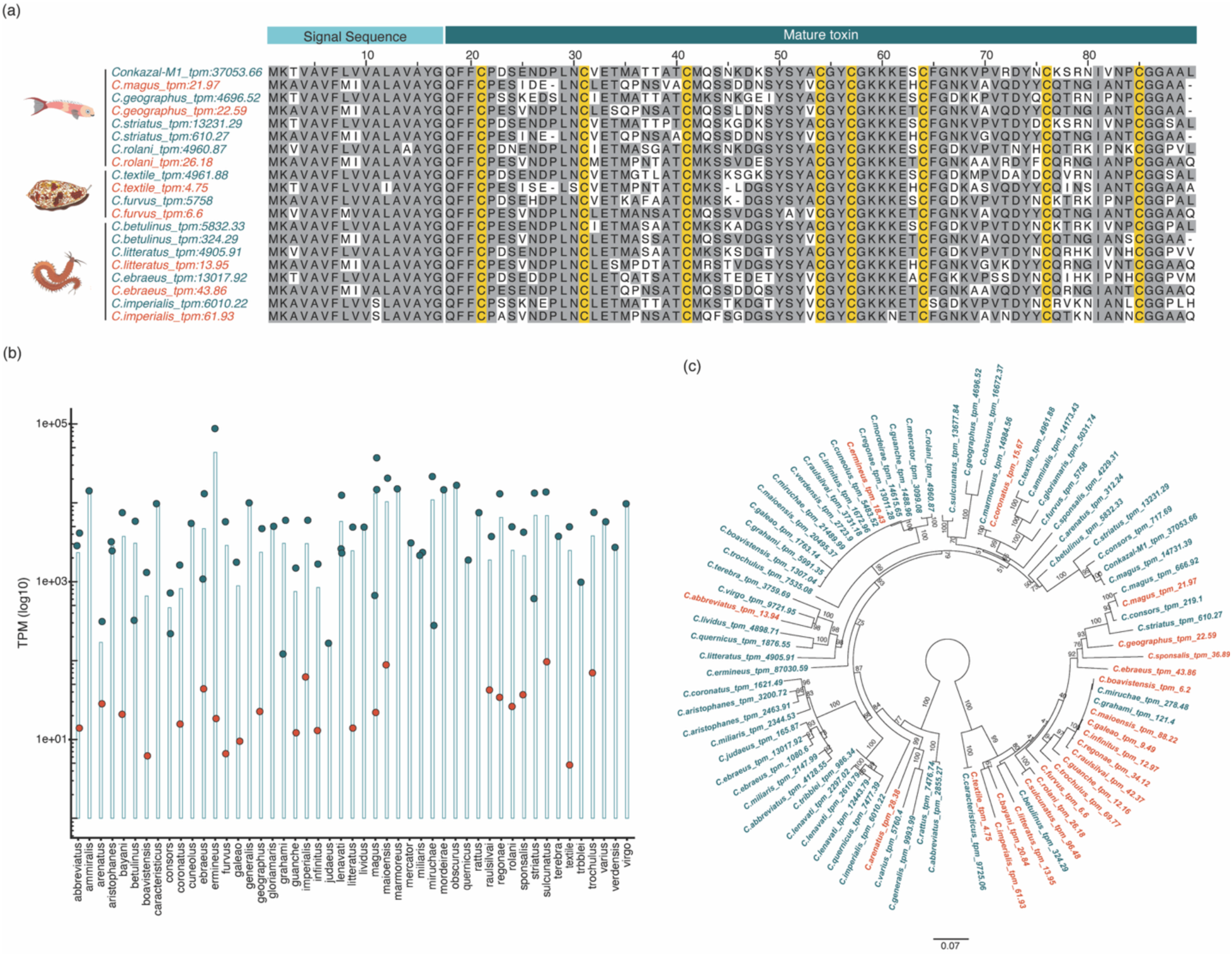
The MKAVAs are encoded by either low- or high-expressing transcripts. **(a)** Multiple sequence alignment of select *Conus* precursor sequences from the MKAVA superfamily. Sequences encoded by high- or low-expressing transcripts are indicated in dark teal and orange, respectively. The signal sequence and mature toxin regions are identified by light and dark teal bars above the alignment. The prey preference of each species (i.e. fish, other molluscs, or worms) is indicated by the illustrations, illustrations, kindly provided by Dr. Paula Florez Salcedo, on the left. **(b)** Gene expression levels of 85 MKAVA transcripts from 49 cone snail species. Dark teal and orange circles indicate high (>100 tpm) and low (<100 tpm) expressors, respectively, with light teal bars showing the average tpm-value of the transcripts for each species. **(c)** Phylogenetic reconstruction of the MKAVA transcripts from (b). Maximum-likelihood was performed using IQ-Tree’s UF bootstrap with 1000 replicates, with best fit model TMP3 + G4 selected by the Bayesian Information Criterion. High-expressing transcripts are shown in dark teal and low-expressing in orange. Some bootstrap values were removed for visualization purposes but are shown in Figure S11. The tree was rooted to separate the two paralogous genes.

In *Conus*, all identified sequences lack an apparent propeptide with the predicted mature toxin sequence (∼73 residues) following immediately after the signal sequence. The toxins exhibit a high level of sequence conservation (Figure 1a) – within the mature toxin sequence, 32 residues are >95% conserved (including all eight cysteines), and approximately 44 positions in an alignment of all identified full-length sequences show ≥75% identity (Figure S1). Selection analyses show that approximately 95% of sites evolved under negative to neutral selection (based on comparisons of the M2 and M1a models and the M8 and M7 models in the Phylogenetic Analysis by Maximum Likelihood (PAML) package (28)), while only three sites (T38, K46 and K48) evolved under weak positive selection with >90% likelihood based on dN/dS ratios (see Supplementary File 2).

Generally, venom gland transcripts that are highly expressed across multiple species likely encode peptides that function as toxins. Analysis of the expression levels of MKAVA transcripts revealed that most cone snail species among those analyzed express two transcripts: one with a high level of expression (100–several thousand transcripts per million (tpm)) and the other with a low level of expression (<100 tpm) (Figure 1b). Phylogenetic analysis of transcripts with high and low expression revealed a well-resolved clustering pattern consistent with their relative expression (Figure 1c). The bipartite nature of the gene tree is reflected at the sequence level, where specific positions distinguish toxins encoded by either high- or low-expressing transcripts (Figure 1a). The high expression of many MKAVA conotoxin-encoding transcripts supports a function of these as toxins. This is further corroborated by their presence in the venom. Using publicly available tandem mass spectrometry data of the venom of *Conus geographus* and *Conus marmoreus*, we identify several tryptic peptides corresponding to MKAVA sequences from these species (Figure S2).

### Conkazal-M1 can be produced in the fully oxidized form with the csDisCoTune *E. coli* expression platform

To investigate the structure and function of the MKAVA superfamily, we selected a high-expressing precursor sequence from the magician cone, *C. magus*, for recombinant expression and characterization. This toxin, the first characterized member of its family, was found to adopt a KPI structure (see below), and was therefore named “conkazal-M1” (where M stands for *magus*). Accordingly, we will use the term “conkazal” rather than MKAVA to refer to this family of conotoxins.

Conkazal-M1 was recombinantly expressed in *E. coli* using the csDisCoTune system (29). This expression platform, a recent further development of the original CyDisCo system (30) and our own csCyDisCo system (31), is specifically tailored for the expression of conotoxins and allows for the production of correctly folded, fully oxidized disulfide-rich proteins in the bacterial cytosol by co-expression of three foldases encoded by an auxiliary plasmid – the Erv1p oxidase, human protein disulfide isomerase (hPDI), and a conotoxin-specific protein disulfide isomerase (csPDI) (32). Conkazal-M1 was expressed as a fusion protein with an N-terminal ubiquitin (Ub) tag containing an internal loop modified to contain ten consecutive histidine residues (His_10_) and a tobacco etch virus protease (TEVp) recognition site, allowing for release of the mature toxin (33).

SDS-PAGE analysis of small-scale test expressions of Ub-His_10_-conkazal-M1 alone, Ub-His_10_-conkazal-M1 with the csDisCoTune-encoded foldases, or with the foldases encoded by the csCyDisCo expression platform clearly showed the beneficial effect of both expression systems on Ub-His_10_-conkazal-M1 production. Both systems resulted in the expression of soluble Ub-His_10_-conkazal-M1 fusion protein, in contrast to expression without the foldases, where equivalent total protein levels were observed but the fusion protein was predominantly found in the insoluble fraction in a reduced and/or partially oxidized state (Figure S3a). Because the csDisCoTune system produced higher levels of oxidized Ub-His_10_-conkazal-M1 compared to the csCyDisCo system, large-scale expression was carried out with csDisCoTune. Cleavage of affinity-purified Ub-His_10_-conkazal-M1 by TEVp resulted in liberation of the 73-residue mature region of conkazal-M1, which was purified (>95% purity) to a final yield of just over 5 mg per liter culture (Figure S3b).

Disulfide-bond formation in the purified toxin was verified by SDS-PAGE analysis, where a clear downward mobility shift was observed upon addition of a reducing agent (Figure S3b), as well as by Q-TOF mass spectrometry, which revealed the presence of two species in the purified preparation. The primary species displayed a monoisotopic mass of 7990.54 Da, in accordance with the theoretical mass (7990.55 Da) of the fully oxidized protein (Figure S3c). The secondary component exhibited a monoisotopic mass of 7973.49 Da, i.e., 17 Da lower than the theoretical mass. We attribute this reduction in mass to modification of the N-terminal Gln to a pyroglutamate, a post-translational modification commonly found in conotoxins (34). This modification can be enzymatically catalyzed by glutaminyl cyclase but is also known to occur spontaneously (35).

### The NMR structure of conkazal-M1 reveals a Kazal-type protease inhibitor fold

The three-dimensional structure of ^15^N and ^13^C stable isotope-labelled conkazal-M1 was solved in solution using distance and torsion angle restraints obtained by NMR spectroscopy. We assigned the chemical shifts of 79% of the backbone ^1^H, ^13^C, and ^15^N resonances and 66% of sidechain resonances. In total, 69% of the chemical shifts were assigned. We observed no resonances from the first seven residues. 602 short distances assigned in NOESY spectra were combined with 100 dihedral angles calculated from chemical shifts and used as restraints in structure calculations (Table 1). The structure is well ordered from residue 10 to the C-terminus with an RMSD value of 0.31Å for the positions of the backbone atoms in the 20 models that represent the structure (Figure S4a).

**Table 1.**
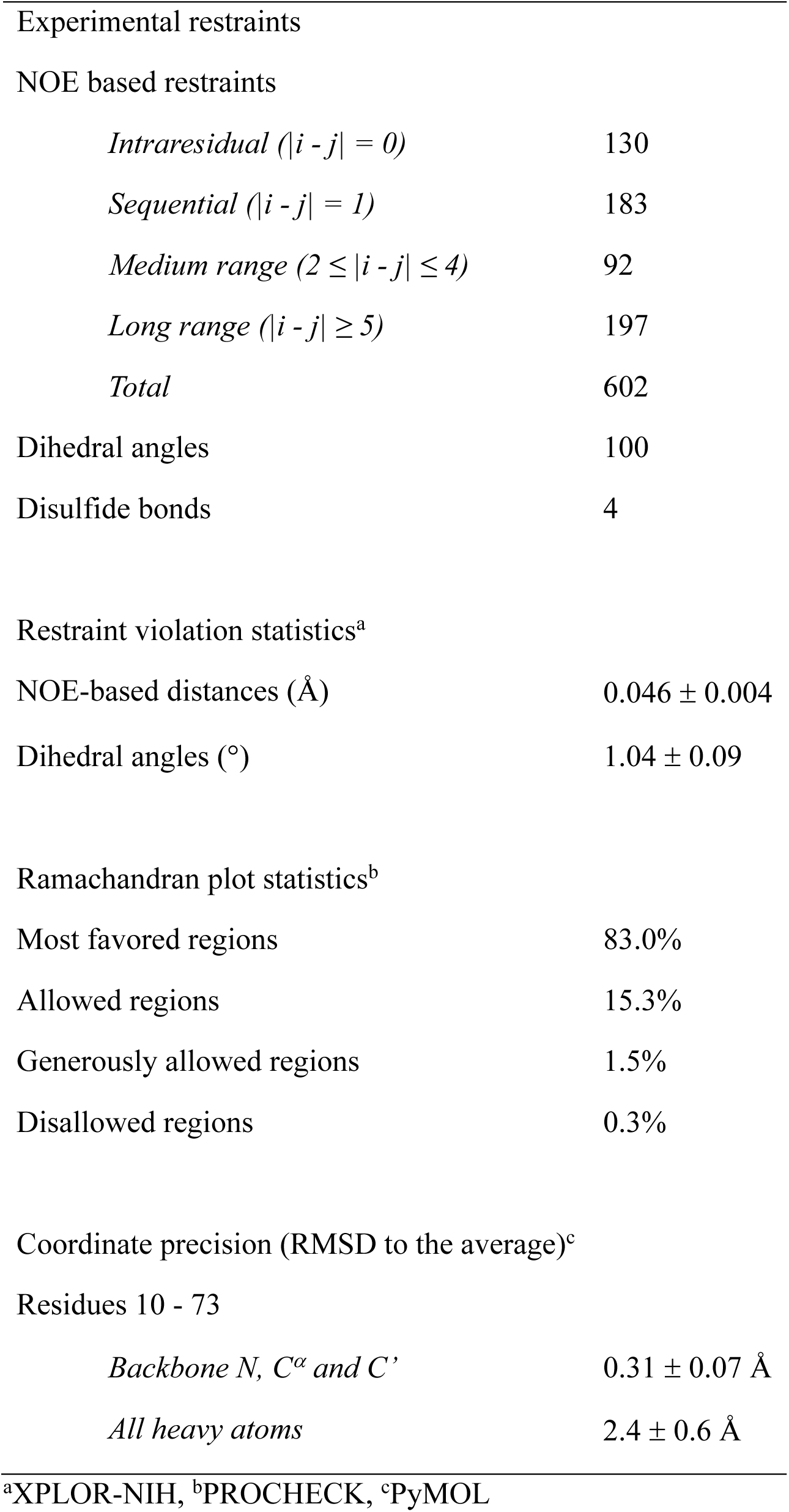
Statistics for the ensemble of 20 conformers representing conkazal-M1.

The structure consists of a central three-stranded antiparallel β-sheet and two α-helices: α-helix 1, which is parallel to the β-strands, and α-helix 2, which is almost perpendicular to α-helix 1 (Figure 2a and b). The N- and C-termini are tethered to the core structure via two disulfide bridges. Located at each end of α-helix 1, Cys40 and Cys48 form disulfide bonds with the N-terminal Cys14 and Cys4, respectively. The C-terminus is affixed to the short loop connecting β-strand 2 and α-helix 1 by the bridge connecting Cys37 and Cys68. The last disulfide bond is formed by Cys24 and Cys59, thus connecting β-strand 1 and α-helix 2.

**Figure 2.**
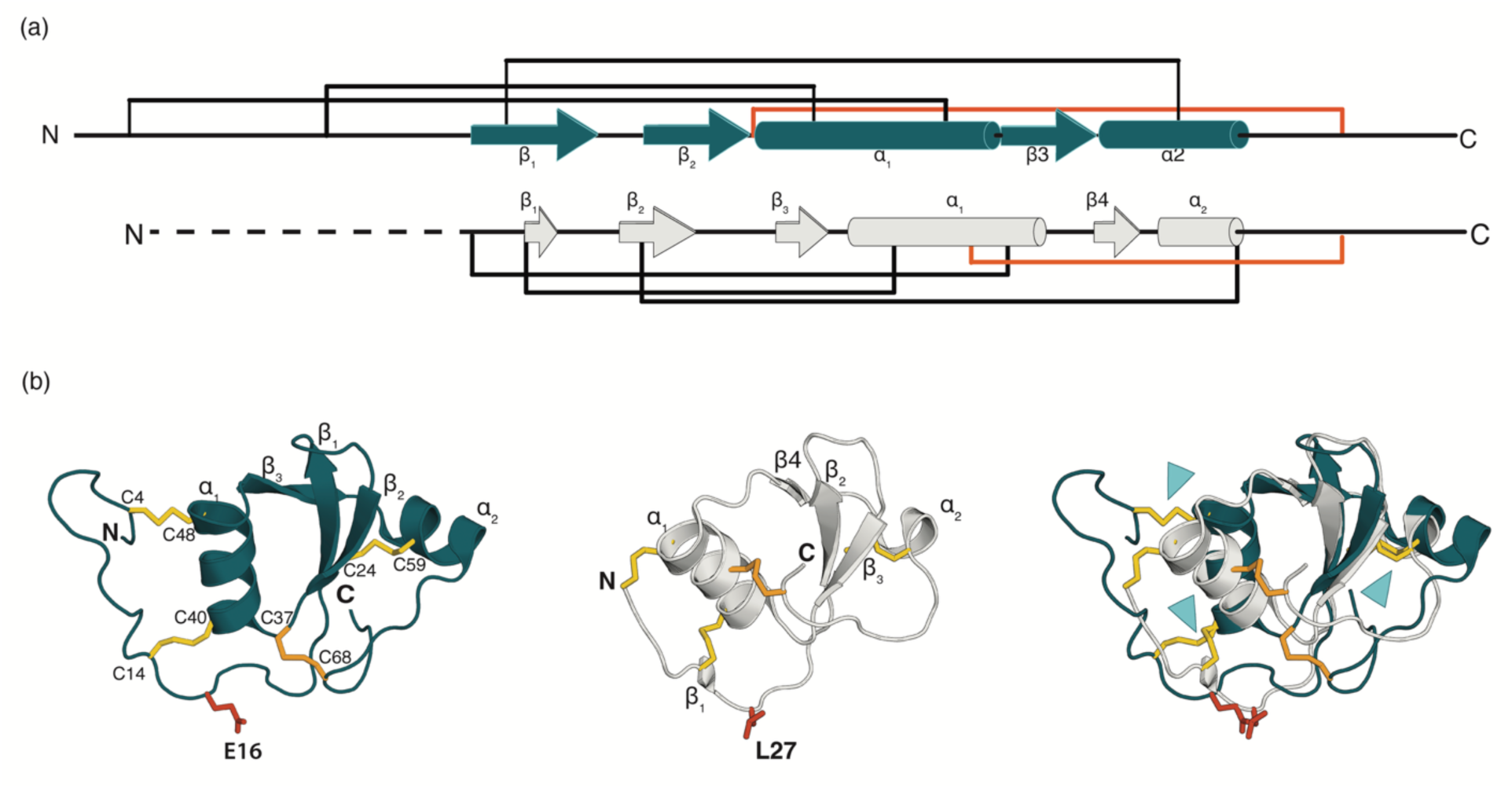
Conkazal-M1 harbors a KPI fold. **(a)**. Graphic representations of Conkazal-M1 (teal) and the Kazal-type protease inhibitor greglin from the locust *Schistocerca gregaria* (gray). Disulfide bridges are represented as brackets. The additional disulfide bridge in conkazal-M1 and greglin (compared to classical KPIs) is shown in orange. The dotted line represents the first 19 residues of greglin not present in the sequence used for X-ray crystallography. **(b)**. Cartoon representation of the conkazal-M1 NMR structure (PDB: 9SLR) (left), cartoon representation of the greglin X-ray crystallography structure (PDB:4GI3) (middle), and superposition of conkazal-M1 and greglin with the three conserved disulfide bridges labeled with blue arrowheads (right). Disulfide bridges are colored yellow or orange in the case of the additional disulfide bride, and the amino acid residue at the P1 position is shown in red.

As structural similarity often correlates with conserved functional properties, we used FoldSeek (36) and PDB eFold (37) to identify proteins structurally similar to conkazal-M1. Both search engines identified several Kazal-type protease inhibitors, such as greglin (38), among the most similar structures to conkazal-M1. Conkazal-M1 and greglin (Figure 2a) superpose with an RMSD value of 2.87Å over 48 residues (Figure 2b). Both structures harbor the KPI core structure as well as an additional C-terminal α-helix not found in classical KPIs. The three characteristic disulfide bridges of the classical KPIs are present in both conkazal-M1 and greglin and similarly positioned in the two structures (blue arrowheads, Figure 2b). Both proteins harbor a fourth disulfide bridge that tethers the C-terminal regions to the core structure of the proteins (Figure 2a; orange). Finally, conkazal-M1 has a longer loop between the first and second cysteine residues.

### Conkazal-M1 inhibits the proteolytic activity of Subtilisin A

Because conkazal-M1 adopts a Kazal fold, we speculated that the toxin may exhibit inhibitory activity against serine proteases. Consistent with this hypothesis, Glu16 at the P1 position in conkazal-M1 is solvent-exposed in all 20 of the lowest energy structures (Figure S4a). Still, we note that although AlphaFold accurately predicts the overall fold of the conkazals, as illustrated for Conkazal-S1 from *C. striatus* (Figure S4b), the Glu side chain at the P1 position points inward in many structural models and is predicted to form hydrogen bonds with the main chain NH groups from Tyr38 and Cys39. This observation informed our decision to determine the NMR structure of the protein and provides an important reminder that AF does not always place side chains accurately (39).

To experimentally determine a potential function of conkazal-M1 as a protease inhibitor, we investigated the proteolytic activities of chymotrypsin, Endoproteinase GluC, and Subtilisin A (SubA) in the presence of increasing amounts (0.25 µM – 3 µM) of conkazal-M1. GluC was chosen as a possible candidate due to its preferential cleavage of peptide bonds C-terminal to glutamic acid residues. SubA is a non-specific protease that favors cleavage after hydrophobic residues but is not restricted to them. For instance, a Glu residue at the P1 position in the second domain of the CrSPI two-domain Kazal-type protein from horseshoe crab is the primary residue for mediating the contacts with subtilisin in the co-crystal of the complex between the two proteins (40).

SubA activity was monitored using the chromogenic substrate N-succinyl-Ala-Ala-Pro-Phe-pNA over a range of substrate concentrations in the presence and absence of conkazal-M1 (Figure 3). Initial rates were estimated from changes in absorbance at 405 nm over time and were converted to reaction rates (µM/min/mg enzyme). Nonlinear regression of the initial rate data was performed using Michaelis-Menten and competitive, non-competitive and uncompetitive inhibition models. Model comparison by Akaike Information Criterion indicated that competitive inhibition best described the data – as expected for a KPI – with estimated kinetic parameters of Km=201 µM (Confidence Interval (C.I.): 173–234 µM), Vmax= 12,888 µM/min/mg (C.I.: 12,357-13,456 µM/min/mg) and Ki=1.1 µM (C.I.: 0.92-1.5 µM) (Figure 3). We performed similar investigations with chymotrypsin and Endoproteinase GluC in the absence and presence (0.5 µM-2 µM) of conkazal-M1, where no effect of the toxin on enzyme activity was discerned (Figure S5).

**Figure 3.**
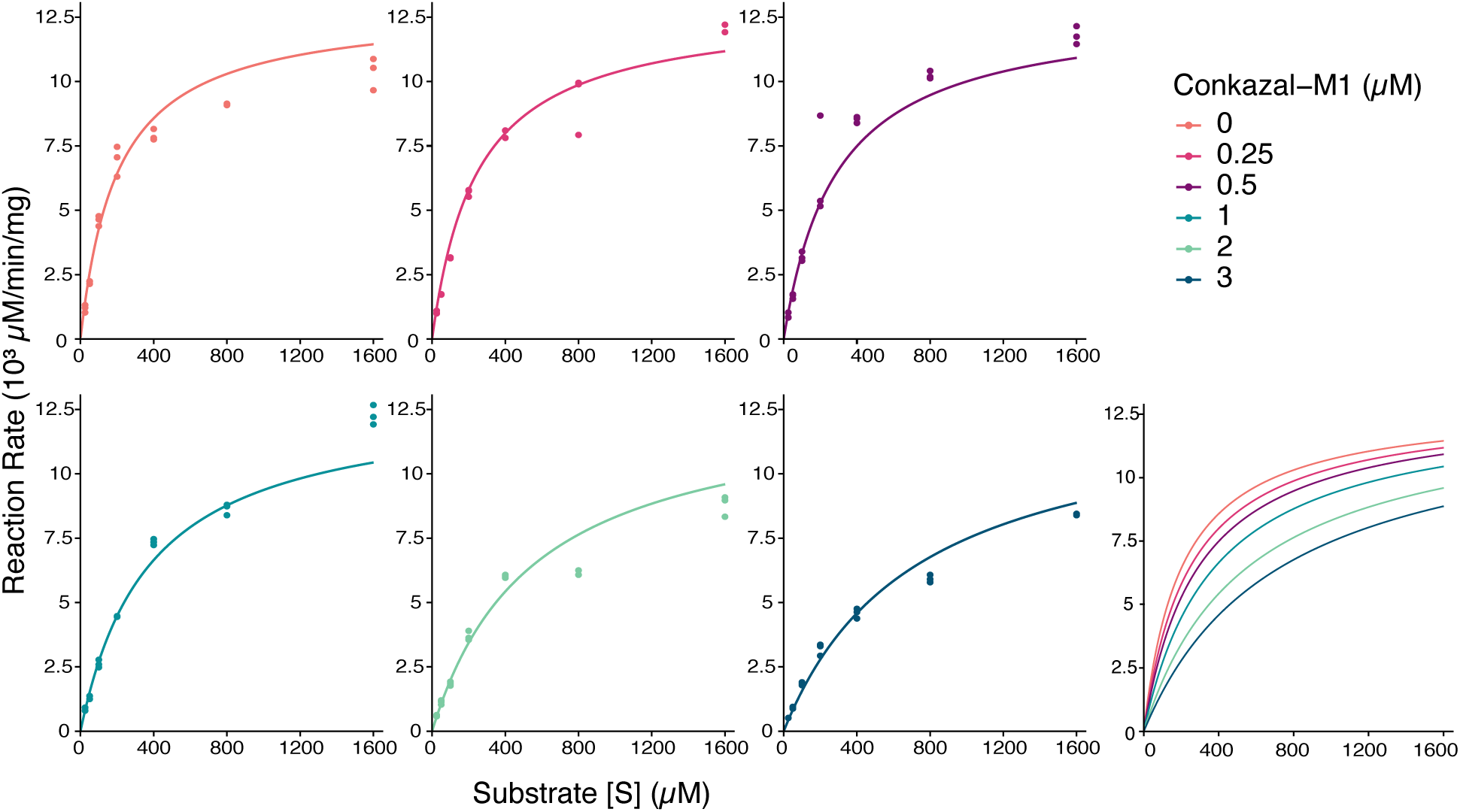
Conkazal-M1 inhibits proteolytic activity of Subtilisin A. Kinetic analysis of Subtilisin A in the presence of increasing concentrations of conkazal-M1. Reaction rates were measured across a range of substrate concentrations and were normalized to enzyme mass. The plots display experimental data points (colored according to inhibitor concentration: 0 μM (orange), 0.25 μM (pink), 0.5 μM (purple), 1 μM (light teal), 2 μM (cyan) and 3 μM (dark teal)) overlaid with lines showing the best-fit model (competitive inhibition). The lower right plot shows all fits overlaid without datapoints to more clearly illustrate the inhibitory effect of conkazal-M1.

### Conkazal-M1 modulates calcium transients in mouse sensory neurons

A wide variety of venom peptides function as neurotoxins that target neuronal ion channels. Therefore, we next assessed the bioactivity of conkazal-M1 using the calcium imaging-based constellation pharmacology assay (Teichert et al, 2012). This assay enables the monitoring of Ca^2+^ influx in primary cell cultures obtained from mouse dorsal root ganglions (DRGs), which comprise various types of sensory neurons, including nociceptors, mechanoreceptors, and proprioceptors, that respond to different sensory modalities. During each experiment, intracellular calcium levels were simultaneously monitored in 500-1,000 cells using the Fura-2-AM dye. Cells were assigned to seven major classes based on their size, isolectin B4 staining (non-peptidergic nociceptors) and calcitonin gene-related peptide (CGRP) expression (peptidergic nociceptors) (Fang et al, 2010). To further characterize neuronal subtypes, a series of pharmacological agents (conotoxin *κ*M-RIIIJ, allyl isothiocyanate [AITC], menthol, and capsaicin) were applied at the end of each experiment (Giacobassi et al, 2020) (Figure 4a).

**Figure 4.**
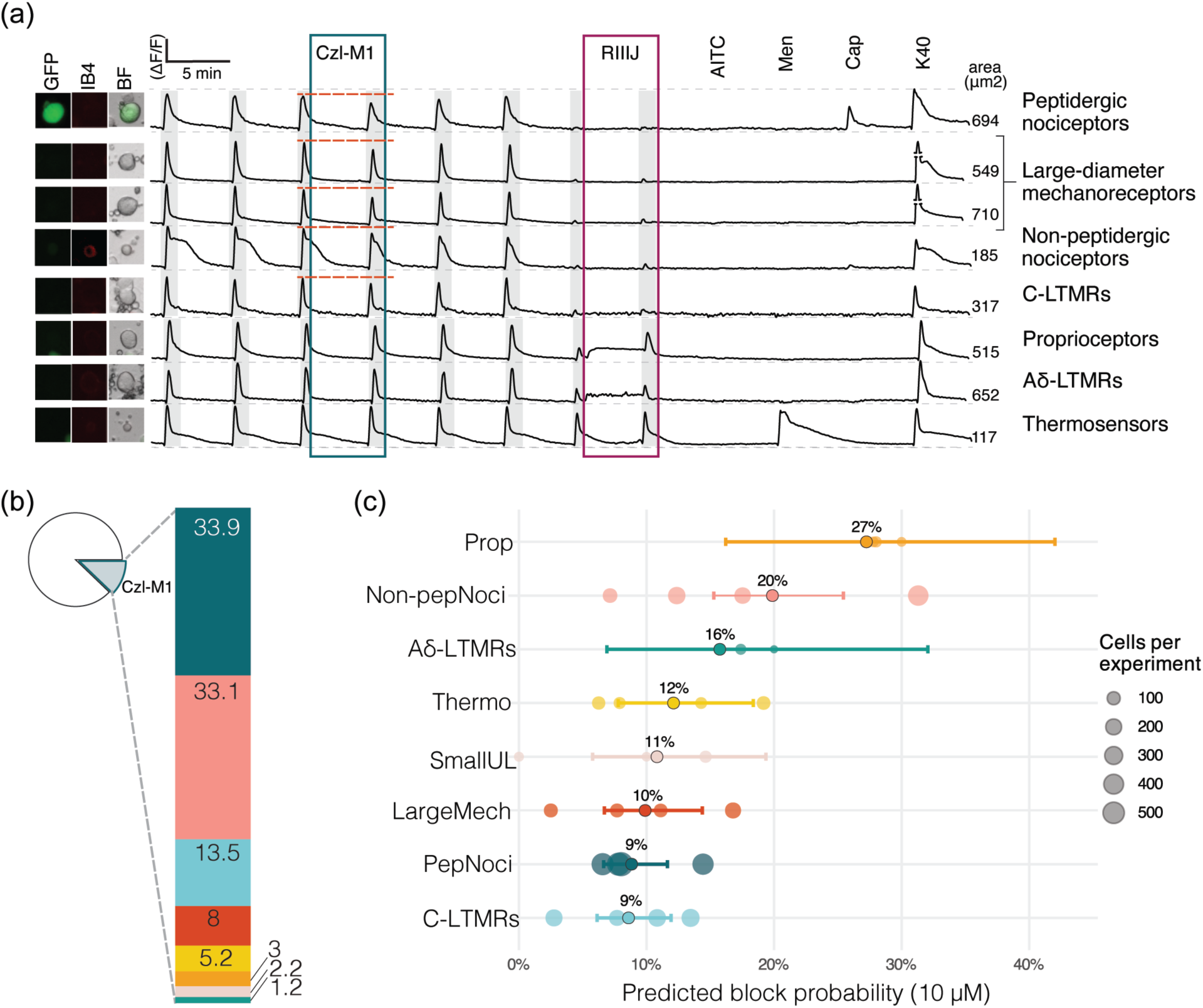
Effects of conkazal-M1 on mouse sensory neurons by constellation pharmacology. (a) Representative calcium-imaging traces of individual neurons responding with a partial block to applications of 10 µM conkazal-M1 (top 5 traces, conkazal-M1 application highlighted by blue box), and neurons that did not respond (bottom 3 traces). Light grey shading indicates KCl (30 mM, except for stimulation prior to RIIIJ treatment, in which 20 mM KCl was used) depolarization pulses. Each trace represents the calcium signal (ΔF/F) of the neuron pictured on the left (GFP: CGRP expression; IB4: isolectin B4 staining; BF: brightfield image; 10x magnification). Pharmacological agents used for cell classification: conotoxin κM-RIIIJ (1 μM, burgundy box); allyl isothiocyanate (AITC, 100 μM), menthol (Men, 400 μM), and capsaicin (Cap, 300 nM). A pulse of 40 mM KCl (K40) was used to assess cell viability at the end of the experiment. Cell size is indicated on the right side of each trace. Aδ-LTMRs: Aδ–low-threshold mechanoreceptors; CGRP: calcitonin-gene related peptide; C-LTMR: C-low threshold mechanoreceptors; GFP: green fluorescent protein; N16: capsaicin-positive small-diameter neurons; RIIIJ: conotoxin *κ*M-RIIIJ. Pie chart representing the population of neurons analyzed (n *=* 4,215) across four independent experiments, highlighting the 12% of conkazal-M1 (10 μM)-sensitive sensory neurons. Bar graph showing the percentage of each class of conkazal-M1-sensitive DRG neurons. Coloration as defined in Panel (c). (**c**) Estimated block probability by neuronal cell type. Each point represents the model-estimated probability that neurons of a given class were blocked by the 10 µM test compound, based on a generalized linear mixed-effects model (logistic link) with experiment as a random intercept. Error bars denote 95% confidence intervals for the marginal means, and translucent circles indicate observed block rates within individual experiments (point size ∝ number of cells). This analysis accounts for between-experiment variability and unequal sample sizes, providing population-level estimates of block probability for each neuronal cell type. The effect of cell type was highly significant (χ²(8) = 276.98, p < 0.0001), indicating robust differences in block probability across neuronal classes. Source data for individual traces and quantifications found in Supplementary File 4. Prop (orange): proprioceptors; Non-pepNoci (pink): non-peptidergic nociceptors; Aδ-LTMRs (green): Aδ–low-threshold mechanoreceptors; Thermo (yellow): Thermosensors; SmallUL (light pink): small-diameter unlabelled neurons; LargeMech: (red) Large-diameter mechanoreceptors; PepNoci (dark teal): Peptidergic nociceptors; C-LTMR (light blue): C-low threshold mechanoreceptors.

Dissociated DRG neurons were depolarized using 15 s pulses of high-potassium extracellular solution (30 mM KCl), resulting in intracellular Ca^2+^ elevation, as indicated by the rise in Fura-2 signal (Figure 4a). At the lowest concentration tested (1 μM), conkazal-M1 did not produce significant effects on any neuronal subclasses. In contrast, application of 10 μM conkazal-M1 reduced the KCl–evoked calcium signals in a subset of cells (12% of the 4,215 cells analyzed) (Figure 4b). The blockade of calcium transients was predominantly observed in small (<400 μm^2^) peptidergic nociceptive neurons (33.9% of all affected cells) (identified by the expression of CGRP genetically labeled with green fluorescent protein) (Figure 4b). Other neuronal subclasses affected by 10 μM conkazal-M1 included large-diameter (>400 μm^2^) mechanoreceptors (8%), a subset of non-peptidergic nociceptors (33.1%), and C-low threshold mechanoreceptors (C-LTMR; 13.5%). Thus, conkazal-M1 inhibits KCl-induced calcium influx in a subset of neuronal cells, albeit with low affinity. To account for the observation that the frequency of neuronal cell types varies considerably among DRG neurons (Figure S6), we quantified the block probability according to cell type (Figure 4c). Most strikingly, considering that proprioceptors constitute only ∼ 3% of all cells (Figure S6), 27% of these were blocked by conkazal-M1 (Figure 4c), indicating that the target of conkazal-M1 is likely most prevalent in this population.

Because conkazal-M1 originates from the venom of *C. magus*, a fish-hunting cone snail, we hypothesized that the toxin might elicit measurable effects in a fish model. To test this, we injected goldfish (*Carassius auratus*) with 5 or 10 nmol of toxin via both intraperitoneal and intramuscular routes (N=3 for each group) and monitored them closely for a period of three hours post-injection. No overt phenotypic effects were observed in either treatment group compared with the controls (data not shown).

### The conkazals are evolutionarily related to schistosomin

To identify potential homologs of conkazals outside Conoidea and thereby gain insight into the evolutionary history of these proteins, we used the conkazal-M1 precursor sequence to perform homology searches against the NCBI non-redundant protein database and transcriptome shotgun assemblies.

The searches revealed sequence similarity, including conservation of all eight cysteines, with schistosomin-like proteins from several molluscs (Figures 5a and S7). Schistosomin is seemingly absent in other phyla than molluscs and was first described in the freshwater pond snail, *Lymnaea stagnalis*, as a peptide upregulated in response to infection with the schistosome parasite *Trichobilharzia ocellata* (41) (see also Discussion). Overall, only modest sequence conservation is observed across the conkazals and schistosomins. In addition to the cysteines, three residues show high conservation. These comprise a glutamate residue located two residues after the second cysteine in both groups of proteins, i.e. at the P1 position of KPIs, at position 34 in the alignment, a proline immediately following the first Cys residue, and a positively charged residue at position 62 in the alignment (Figure 5a and S7).

**Figure 5.**
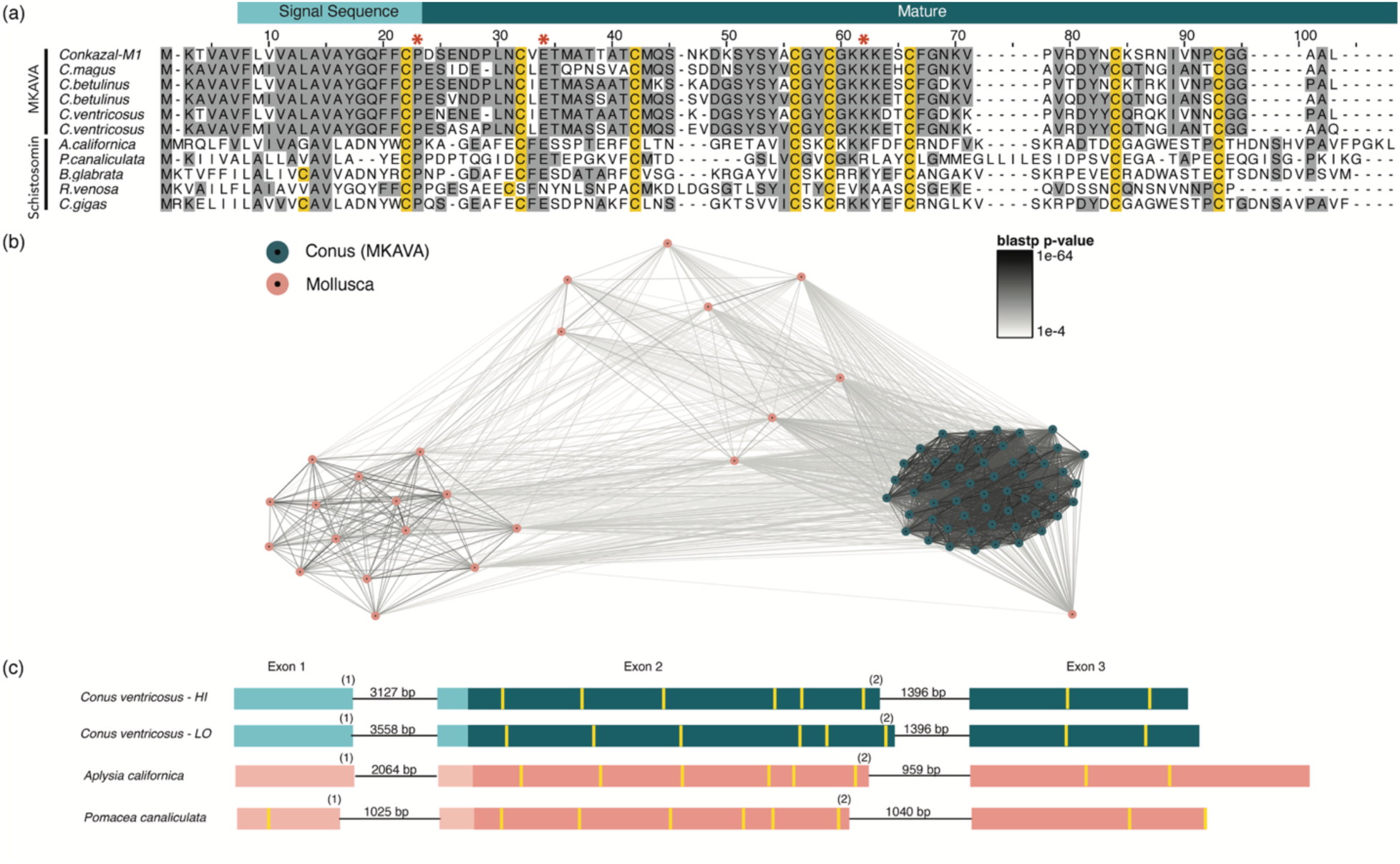
The MKAVAs are evolutionarily related to schistosomin. **(a)** Multiple sequence alignment of MKAVAs and schistosomin-like sequences from *Aplysia californica* (XP_005098364.1)*, Pomacea canaliculata* (PVD28780.1)*, Biomphalaria glabrata* (NP_001298209.1)*, Rapana venosa* (GDIA01100366.1) and *Crassostrea gigas* (GHAT01224603.1). Red asterisks denote amino acid residues within the mature protein region exhibiting over 88% identity across all sequences analyzed (see Supplementary Figure S7). **(b)** CLANS clustering analysis of conotoxins and schistosomins. Deep teal: conotoxins from MKAVA superfamily; pink: molluscan schistosomin. Nodes depict individual precursor sequences, and the edges correspond to BLOSUM62 blastp p-values. **(c)** Gene structure analysis of toxins with high or low expression from *Conus ventricosus*, and of schistosomins from *A. californica* and *P. canaticulata*. The exons are depicted as boxes proportional to the length of the sequences, while introns are represented as thin lines (with sequence length labelled). The intron phases (0, 1, or 2) are indicated prior to the introns. The yellow lines indicate cysteine positions within the exons.

A CLANS (CLuster ANalysis of Sequences) clustering analysis (42), which groups protein families based on all-against-all pairwise protein sequence similarities, revealed a clear similarity of the *Conus-*derived conkazal sequences with molluscan schistosomins from gastropods (Figure 5b; all sequences used in this analysis are provided in Supplementary File 3). Moreover, gene structure analysis revealed a common gene structure of conkazals and molluscan schistosomins. The genes have a common phase-1 intron located immediately downstream of the signal sequence and a conserved phase-2 intron located between the sixth and seventh cysteine residues (Figure 5c). The common gene structures further support an evolutionary relationship between conkazal conotoxins and schistosomins.

Given the conservation of all cysteine residues and the common evolutionary origin, it was not surprising that the AlphaFold-predicted structure of a schistosomin from *Lymnaea stagnalis* was found as the top hit in the FoldSeek search with conkazal-M1. We also identified an unpublished crystal structure of schistosomin from another freshwater snail, *Biomphalaria glabrata* (PDB ID: 9FDO), in the PDB. Conkazal-M1 and this schistosomin (Figure S8a) superpose with an RMSD value of 1.9 Å over 38 Cα-atoms (Figure S8b). Both structures adopt the KPI core structure as well as an additional C-terminal α-helix not found in classical KPIs. The three characteristic disulfide bridges of the classical KPIs are present in both conkazal-M1 and schistosomin and are similarly positioned in the two structures (light blue arrowheads, Figure S8b). Both proteins harbor a fourth disulfide bridge that tethers the C-terminal regions to the core structure of the proteins (Figure S8a).

While molluscan species exhibit a high degree of intrachromosomal gene translocation, macrosynteny tends to be conserved. We therefore compared the genomic location of conkazal genes from *C. ventricosus* with schistosomins from representative gastropods (Figure S9). The two conkazal genes in *C. ventricosus* are adjacent to each other on scaffold 23, which represents one of the chromosome-scale scaffolds in the cone snail genome. This chromosome shows homology to chromosome NC_037594.1 of the gastropod *Pomacea canaliculata* and chromosome NC_074711.1 of another gastropod, *Biomphalaria glabrata* (Figure S9, left). However, neither of these chromosomes contains the schistosomin genes, which are located on NC_037595.1 and NC_074716.1, respectively. Surprisingly, both *P. canaliculata* and *B. glabrata* have multiple schistosomin gene copies located in tandem, though the neighboring genes show no syntenic relationship (Figure S9, right). Overall, while conkazals show a high level of similarity to schistosomins, strongly suggesting an evolutionary relationship, the genes are not part of a common gene linkage group.

## DISCUSSION

Here, we provide the first characterization of a toxin from the conkazal conotoxin family, conkazal-M1. The conkazals constitute a large and widespread toxin family found across phylogenetically diverse *Conus* species (from all three feeding groups) and several other species in the Conoidean superfamily, indicating an important functional role of these peptides. The common evolutionary history with the schistosomins indicates that an ancestral gene was recruited into the venom in early Conoidea and then acquired neuroactive functions. However, given that the function of schistosomins remains unclear, the exact process of recruitment is presently unknown.

Many conkazal transcripts are highly expressed in the venom gland, and we confirmed their presence in venom by proteomic analysis, supporting a role as toxins. We show that conkazal-M1 can perform dual functions as a protease inhibitor and as a partial inhibitor of KCl-stimulated calcium influx in sensory neurons. Such dual activities have also been reported in Kunitz-type venom proteins (11–14). In the context of constellation pharmacology, the partial suppression of KCl-evoked calcium transients by conkazal-M1 provides preliminary evidence that this peptide can modulate ion channel activity (directly or indirectly) in sensory neurons. The modest inhibition at 10 µM indicates either a low-affinity or indirect mechanism, potentially consistent with dual-function peptides in which ion-channel modulation is a secondary activity to protease inhibition, as further discussed below. Future studies are needed to identify the target and mode of action of conkazal-M1 in neuronal cells.

In the context of a protease inhibitor function, subtilisin-targeting KPIs with a Glu at the P1 position have previously been characterized (see e.g., (40, 43–45)). It is therefore unsurprising that conkazal-M1, with its Glu16 at the P1 position, was able to inhibit subtilisin (although additional residues in the reactive-site loop likely also contribute to substrate specificity). While certain venoms appear to harbor microbiomes (46), bacterial subtilisin is an unlikely physiological target of the protease-inhibitory function of conkazal-M1. This is also reflected in the relatively high *K*_i_ value of conkazal-M1 for subtilisin when compared to many other subtilisin inhibitors, such as greglin. In principle, as a protease inhibitor, conkazal-M1 could function either in the venom, to regulate venom protease activity and maintain stability before delivery, or in the prey, to exert a deleterious effect upon delivery. At present, we cannot distinguish between these possibilities, although we note that we have found no evidence in the literature for an endogenous regulatory function in the venom of protease inhibitors in other venomous organisms.

When comparing conkazal sequences with the schistosomins, the Glu residue at the P1 position is one of the few residues conserved across species (Figure 5). This observation supports the hypothesis that the conkazals and mollusc schistosomins function as KPIs. However, Glu33 in the crystal structure of schistosomin is not solvent-exposed (as would be expected of a P1 residue) but points towards the interior of the structure (Figure S8b). Overall, the potential function of schistosomin as a KPI remains unresolved. In addition to the eight cysteine residues, only two other residues are as highly conserved in conkazals and schistosomins as the P1 Glu: a proline residue immediately following the first Cys residue, likely playing a structural role, and a surface-exposed lysine residue at a position corresponding to Lys43 in the mature conkazal-M1 sequence, which is likely to have a functional role.

Schistosomiasis, a parasitic infection, is caused by flatworms known as schistosomes. In the schistosomiasis life cycle, freshwater snails serve as intermediate hosts that release infectious larvae into the water. These larvae can penetrate human skin, gaining access to the host, where the subsequent release of eggs can cause a range of symptoms, including serious damage to organs such as the liver and bladder. In the snail-schistosome system *L. stagnalis*-*T. ocellata*, schistosomin was initially proposed to be a neuropeptide induced by parasite infection, but subsequent studies showed that it is produced in connective tissue cells and in hemocytes (reviewed in (47)). In the hemolymph, schistosomin was proposed to act as an antagonist of gonadotropic hormones, thereby inhibiting reproductive activity in snails. However, in the *Biomphalaria glabrata*-*Schistosoma mansoni* system, no correlation was found between parasite infection and schistosomin levels and infection did not result in parasitic castration (48). Thus, the physiological function of schistosomin remains unclear, and further investigation is warranted, particularly with its potential function as a KPI in mind.

An ancestral protease inhibitor that acquires the ability to modulate neuronal activity while retaining protease inhibition represents a functional change in which purifying selection preserves the original function, while weak-to-moderate selection can favor the emergence of a novel activity (6, 49). In contrast, a complete shift toward modulation of neuronal activity accompanied by partial or total loss of protease activity can be explained by strong positive selection for the new function, coupled with relaxed constraint on the ancestral activity. Because we detect no evidence of strong positive selection, we favor the first evolutionary model. In this scenario, conkazals would be “moonlighting” proteins, facing an adaptive conflict when mutations that enhance one activity compromise the other. Such conflicts are typically resolved through gene duplication, followed by subfunctionalization, which partitions the ancestral roles between paralogs, or through neofunctionalization of one copy. Our data do not support a neofunctionalization event following duplication. If such an event had occurred, we would expect to detect both an ancestral, schistosomin-like gene and a duplicated conkazal gene. Instead, we only observe conkazal genes when interrogating available cone snail genome data, consistent with a moonlighting role. This interpretation is further supported by the expression of conkazals in tissues other than the venom gland, including the venom bulb, nerve ring, and foot (see Materials and Methods section for SRA entry numbers), where these peptides likely play an endophysiological function.

We note that multi-domain Kazal-type proteins are found in many venomous animals, including the two-domain Kazal-type toxins BQTX from shrews (19) and Rhodniin from the kissing bug *Rhodnius prolixus* (17, 50). In general, secreted cysteine-rich repeat-proteins (51) – including a broad range of venom peptides – are common in nature and have been cataloged in the ScrepYard database (52). Their multi-domain structure enables multivalent substrate interactions, allowing them to bind targets strongly through an avidity effect. Indeed, Rhodniin binds its substrate thrombin through multivalent interactions involving the reactive-site loop of one domain and electrostatic interactions of the second domain, together conferring a picomolar affinity (17, 50). Analogous to the two-domain Kazal-type toxins described above, we have recently identified a family of such conotoxins, the MKTAV family. As shown in Figure S10a-b, both Kazal domains of these conotoxins, with few exceptions, contain a Glu or Asp residue three positions C-terminal to the second Cys in the mature sequence. Although not located at the canonical P1 position, we speculate that this residue could still perform the inhibitory function of a P1 residue. The AlphaFold 3 predictions of the MKTAVs reveal two very similar Kazal-type domains connected by a short linker, with each domain closely resembling the structure of conkazal-M1 (Figure S10c-e).

In this study, we identify two previously unrecognized families of conotoxins, conkazal and MKTAV, which encode single- and two-domain Kazal-type structures, respectively. We uncover the dual function of conkazal-M1, acting both as a neuroactive peptide with the ability to inhibit calcium transients in a subset of DRGs and as a protease inhibitor. Together, these observations highlight the structural and functional diversity of Kazal-type toxins in *Conus*, extend the current knowledge about the toxin complement available to venomous animals, and illustrate the adaptive conflict that can shape the evolutionary trajectories of dual function toxins.

## MATERIALS AND METHODS

### Sequence mining and analysis

Multiple sequence alignments were generated with the MAFFT version 7 (53) online interface using the G-INSi method (54). Alignments were visualized in JalView (55). Schistosomin sequences were obtained using web version of blast to query against the non-redundant (nr) protein database and transcriptome shotgun assemblies with standard settings. Conotoxin sequences were obtained from (25). These sequences were used as input for CLANS (42), which is a sequence-based clustering program that assigns attractive values between sequences based on their BLOSUM62 blastp p-value. In a uniformly repelling force field this causes similar sequences to cluster together. Gene structure analyses were performed with using the protein2genome (p2g) model in Exonerate (v.2.76.2) (56). Macrosynteny analysis was performed with macrosyntR using OrthoFinder to assign orthologous proteins across *C. ventricosus* (GCA_018398815.2)*, P. canaliculata* (GCF_003073045.1), and *B. glabrata* (GCF_947242115.1). The local gene neighborhoods were assessed and visualized in Figure S9. Maximum likelihood gene trees were constructed using alignments generated with MAFFT v.7.526 and the tree was constructed using IQ-TREE v. 2.2.2.3 (57) with UF bootstrap (58) calculation for 1000 replicates. The best model according to the Bayesian Information Criterion was calculated to be TMP3 + G4. The consensus tree was visualized using FigTree (59). Conkazal-like sequences were identified in the NCBI SRA database using the conkazal-M1 amino acid sequence as a query sequence in tblastn searches that retrieved the following SRA entries: SRX8002177, SRR11423812 (Drillidae), SRR26897535, SRR26897526 (Turridae), SRR10192879, SRR10192877 (Profundiconus), SRR2060989 (Terebridae), SRR16493587 (venom bulb), SRR16493599, SRR16493598 (nerve ring), and SRR16493591 (foot).

### Proteomics-based identification of MKAVA peptides in venom

Proteomics files of reduced, alkylated and trypsin digested venom of *Conus geographus* (accession: PXD052869) and *Conus marmoreus* (PXD038992) were retrieved from the PRIDE ProteomeXChange repository and searched for MKAVA sequences using the Byonic software (version 3.2.0 Protein metrics) with the following settings: cleavage site: RK, missed cleavage: 1, Modifications: carbamidomethyl (C, +57.02146), Oxidation (M,P, +15.994915), pyro-Glu (Q, +17.026549), Carboxy (E, +43.989829). High scoring peptide hits (score> 400) were manually verified.

### Plasmid generation

The codon-optimized sequence encoding mature conkazal-M1 was synthesized and inserted (TWIST Bioscience) into the pET39_Ub19 expression vector (33). The resulting plasmid (pLE657) encodes ubiquitin (Ub)-His_10_-tagged conkazal-M1 (Ub-His_10_-conkazal-M1) with a recognition site for TEV protease located immediately N-terminally to the mature toxin sequence.

### Protein expression and preparation of lysate

Chemically competent BL21(DE3) were co-transformed with pLE657 and the csDisCoTune expression plasmid (pLE900) or, for comparison, alone or together with the csCyDisCo plasmid (pLE577). Cells were plated on lysogeny broth (LB) agar supplemented with kanamycin (50 μg/mL) and – when co-transforming with pLE900/pLE577 – chloramphenicol (30 μg/mL). LB medium containing the same type and concentration of antibiotic as in the transformation plates was inoculated with a single colony and incubated for at 37° C for 18 hours at 190 rpm on an orbital shaker.

Initial small-scale expression tests were performed in 50 mL LB medium containing appropriate antibiotics and supplemented with 0.05% glucose. Test cultures were inoculated with 2% overnight culture and grown at 37°C, 190 rpm until OD_600_ reached ∼0.8. Expression was induced by addition of isopropyl-D-1-thiogalactopyranoside (IPTG) to a final concentration of 1 mM and the cultures grown for 18 hours at 25° C with shaking at 190 rpm to allow protein expression. Large-scale expression was performed similarly in 1 L LB medium. Expression performed in 0.5 L M9 minimal medium for stable isotope labelling was performed as described in (60).

Induced cultures were harvested by centrifugation at 4,000 x g for 20 minutes at 4° C. The cell pellets were resuspended in 20 mL lysis buffer (50 mM Tris (pH 8.0), 300 mM NaCl, 20 mM imidazole) per liter culture. Cell lysis was performed using a UP200S ultrasonic processor (Hielscher) while keeping the cells on ice. The cells were lysed with 8 x 30-second pulses at 90% power with 30-second rests between each pulse. Cell debris was subsequently pelleted by centrifugation at 30,000 x g for 45 minutes at 4° C. The cleared supernatant was reserved, and cell pellets were resuspended in an equal volume lysis buffer containing 8 M urea for SDS-PAGE analysis.

### Protein purification

Ub-His_10_-conkazal-M1 was affinity purified from the clarified lysate on an ÄKTA START system equipped with a 5-mL prepacked HisTrap HP (Cytiva) column equilibrated in lysis buffer. The lysate was applied to the column and washed with 30 column volumes (CVs) of lysis buffer before elution of Ub-His_10_-conkazal-M1 with a gradient of 0% to 100% elution buffer (50 mM Tris (pH 8.0), 300 mM NaCl, 400 mM imidazole) developed over 20 CVs. Pooled fractions were dialyzed twice against 2 L cleavage buffer (50 mM Tris (pH 8.0), 300 mM NaCl).

The Ub-His10-conkazal-M1 fusion protein was cleaved using His_6_-tagged TEVp, expressed, and purified essentially as described previously (27). A molar ratio Ub-His_10_-conkazal-M1: His_6_-TEVp of 1:2 was used. To avoid reducing the disulfides in conkazal-M1, His_6_-TEVp – preactivated with 2 mM dithiothreitol (DTT) for 30 minutes at room temperature – was diluted to approximately 0.002 mM DTT by 3 rounds of dilution/concentration in an Amicon Ultra 15 mL 10K Centrifugal Filter (Merck Millipore). TEVp cleavage was performed overnight at room temperature.

To remove uncleaved Ub-His_10_-conkazal-M1, the Ub-His_10_ tag, and His_6_-TEVp, the cleavage mixture was applied to a 5 mL prepacked HisTrapHP (Cytiva) affixed an ÄKTA START equilibrated in cleavage buffer. The flow-through and the first wash fraction were collected. The presence of cleaved conkazal-M1 in the flow-through and wash fractions was investigated by analysis on a 15% SDS-PAGE gel, and the protein-containing fractions pooled. Due to minor remaining impurities, the cleaved conkazal-M1 was subjected to size exclusion chromatography on a Superdex 75 Increase 10/300 GL column (Cytiva) equilibrated in 50 mM Tris (pH 7.8), 150 mM NaCl. Fractions containing purified conkazal-M1 were pooled and stored at 4 C.

### SDS-PAGE analysis

Samples from bacterial expression and subsequent purification steps were separated on 15% glycine SDS-PAGE gels and where indicated samples were treated with 40 mM DTT. Protein bands were visualized with Coomassie brilliant blue stain and images were captured with a BioRad Chemidoc Imaging System.

### Protein concentration determination

Concentration was determined by measuring absorbance at 280 nm and using the theoretical extinction coefficient provided by the Expasy ProtParam tool available through the Expasy bioinformatics resource web portal (61).

### Mass spectrometry

Mass spectrometry was performed on a Waters Xevo G3 QTof system equipped with a ACQUITY UPLC C4 reversed-phase liquid chromatography column (300 Å, 1.7 µm, 2.1 mm × 50 mm) using a sample of purified conkazal-M1 at 100 µM adjusted to pH 2 with trifluoroacetic acid as input.

### NMR spectroscopy

A sample of 500 µM ^13^C,^15^N-labelled conkazal-M1 was prepared in 10 mM 2-(N-morpholino)ethanesulfonic acid (MES), 200 mM NaCl, 5% D_2_O, pH 6.0. For backbone chemical shift assignments ^15^N-HSQC, HNCA, HN(CO)CA, CBCA(CO)NH, CBCANH, CBCA(CO)NH, HNCO, and HN(CA)CO spectra were recorded on a Bruker Avance III HD spectrometer operating at 750 MHz with a triple resonance cryoprobe. Side chain assignments were achieved from HCCONH and HCCH-TOCSY spectra. Distance restraints were obtained from ^15^N-NOESY-HSQC and ^13^C-NOESY-HSQC experiments recorded using a mixing time of 120 ms. The triple resonance spectra were recorded with non-uniform sampling at 25% and reconstructed with qMDD (62). All spectra were processed with nmrPipe (63) and analyzed in CCPNMR analysis (64).

### NMR structure calculations

Automated NOE assignment was performed using Cyana (65). The NOE list was manually refined. XPLOR-NIH (66) was used for the final structural refinement including a torsion angle database potential (67) and an implicit solvent mode (68). For the final ensemble representing the solution structure of conkazal-M1, the 20 structures with the lowest energy were chosen from 100 calculated structures. The Ramachandran-plot statistics was calculated using PROCHECK (69) and the coordinate precisions were calculated by PyMOL (DeLano Scientific). Structure visualization was also performed with PyMOL. The structure has been deposited in the Protein Data Bank (PDB, wwpdb.org) with PDB code 9SLR and chemical shifts and peaks lists from the NOESY spectra have been deposited in the Biological Magnetic Resonance Bank (http://www.bmrb.wisc.edu/) with ID 53313.

### Structural homology search

Structures similar to conkazal-M1 were identified using FoldSeek (36), and structural superpositions were executed manually in PyMOL (DeLano Scientific) utilizing the CEalign command.

### Protease assays

Endoproteinase GluC, bovine pancreas α-chymotrypsin and subtilisin A and substrates for chymotrypsin (N-Succinyl-Gly-Gly-Phe-*p*-nitroanilide) and subtilisin A (N-succinyl-Ala-Ala-Pro-Phe-p-nitroanilide) were all purchased from Merck. Substrate for GluC (Z-Phe-Leu-Glu-pNA) was purchased from Bachem.

Inhibition assays with chymotrypsin and GluC were performed in assay buffer (50 mM Tris, 2 mM CaCl_2_, pH 7.8). Subtilisin A inhibition experiments were performed in assay buffer, pH 8 supplemented with 5 mM CaCl_2_. The proteases (3.6 nM SubA, 100 nM GluC and 400 nM chymotrypsin) were incubated with varying concentrations of conkazal-M1 for 10 min at 29° C. Proteolytic reactions were initiated by addition of the substrates, and absorbance at 405 nm was recorded every 30 seconds for 30 minutes. The 96-well plate was shaken orbitally for 10 seconds immediately prior to measurement. Initial reaction rates were estimated with the web-based tool, IceKat (70). The absorbance-based rates for Subtilisin A were converted to reaction rates in µM/min/mg using the extinction coefficient for AAPF-p-nitroaniline (ε = 8800 M⁻¹cm⁻¹) and an estimated path length of 0.2 cm for 100 μL reaction volumes in 96-well plates. The absorbance-based rates (Abs/min) for Chymotrypsin and GluC were plotted without conversion. The data were fit to four models of enzyme inhibition: Michaelis-Menten (no inhibitor), competitive, uncompetitive, and noncompetitive inhibition.

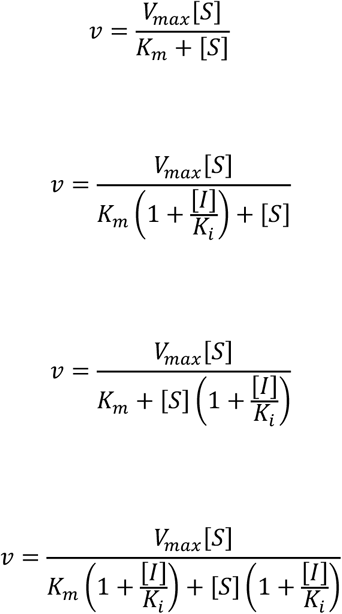

Model comparison using Akaike Information Criterion identified the best fit to the data.

### Cell culture and constellation pharmacology

Experiments were performed as previously described (71). Briefly, all lumbar DRG neurons from C57BL/6 mice 30-40 days old of both genders were dissociated by trypsin digestion and mechanical trituration and plated on polylysine-coated plates. The cells were incubated overnight in a 5% CO_2_ incubator at 37 °C in neuronal culture medium [minimal essential media supplemented with 10% (vol/vol) fetal bovine serum, penicillin (100 U/mL), streptomycin (100 μg/mL), 1× Glutamax, 10 mM HEPES, and 0.4% (wt/vol) glucose, adjusted to pH 7.4]. One hour before calcium imaging experiments, cells were loaded with 2.5 μM fura-2-acetoxymethyl ester (Fura-2-AM, Sigma-Aldrich) and incubated at 37 °C. Cells loaded with Fura-2-AM dye were excited intermittently with 340- and 380-nm light; fluorescence emission was monitored at 510 nm. An image was captured at each excitation wavelength, and the ratio of fluorescence intensities (340/380 nm) was acquired every 2 s to monitor the relative changes in intracellular calcium concentration in each cell as a function of time. Typically, 500-1,000 neurons were imaged for each experiment. The calcium transients were elicited by 15 s applications of depolarizing stimulus (30 mM KCl, except for stimulation prior to RIIIJ treatment, in which 20 mM KCl was used), followed by four washes with extracellular solution [145 mM NaCl, 5 mM KCl, 2 mM CaCl2, 1 mM MgCl2, 1 mM sodium citrate, 10 mM HEPES, and 10 mM glucose, adjusted to pH 7.4]. For activity screening, conkazal-M1 was applied in extracellular solution at 1 and 10 μM between KCl pulses and activity was assessed by significant change in the KCl response after incubation with conkazal-M1. Four different pharmacological agents were used for cell classification: conotoxin *κ*M-RIIIJ (1 μM), AITC (100 μM), menthol (400 μM) and capsaicin (300 nM). At the end of the experiment, cells were incubated with 0.7 mL of Hoechst stain (1000 μg/mL) and Alexa-Fluor 647 Isolectin B4 (2.5 μg/mL) for 5 min. The data was acquired using NIS-Elements and further processed with CellProfiler v.3.1.9 (Jones et al, 2008). Custom-built scripts in Python v.3.7.2 and R were used for further data analysis and visualization. The experimental methods were approved by the Institutional Animal Care and Use Committee (IACUC) of the University of Utah (Protocol number: 17–05017).

### Statistical analysis of calcium imaging data

Each of the four independent experiments included a series of KCl-induced responses. Their magnitude was measured as the maximum area under the curve over any 15-s interval. The deviation of this response after incubation with conkazal-M1 was used to estimate the effect of conkazal-M1. We estimated this effect on each neuron using the linear model (lm function in R) with the following:

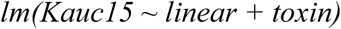

where Kauc15 was the maximum area under the curve for any 15 s interval for each KCl pulse, linear was the sequential trend coded as sequential integers, toxin was an indicator variable for the pulse immediately following incubation with conkazal-M1.

To control for false positives and set appropriate thresholds for significance we used Monte Carlo simulations. Each simulation generated random numbers drawn from a normal distribution with mean and standard deviations of the actual data. The T-stats were estimated for all cells and recorded. This is considered a null distribution. The T-stat estimates from 100 Monte Carlo simulations were used to establish the thresholds for single test cell significance as well as per-experiment significance.

Block responses were analyzed at the level of individual neurons using a generalized linear mixed-effects model (GLMM) implemented in R (version 4.x) with the lme4 and emmeans packages. For each cell, the binary outcome variable block10 μM (1 = blocked, 0 = unblocked) was modeled as a Bernoulli trial with a logit link, using the following structure:

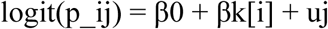

where p_ij is the probability that cell i in experiment j was blocked, βk[i] is the fixed effect associated with its neuronal class, and uj ∼ N(0, σ²_exp) is a random intercept capturing between-experiment variation. This formulation accounts for unequal numbers of cells across experiments and for correlations within experiments. Model fitting used the bobyqa optimizer with 200,000 iterations. Marginal means (estimated probabilities of block for each neuronal class, averaged across experiments) were obtained from the fitted model using emmeans with type = ‘response’, which returns predictions on the probability scale. The overall effect of neuronal cell type on block probability was evaluated using a Wald chi-square test of the fixed effect in the GLMM. This test indicated a highly significant difference among cell types (χ²(8) = 276.98, p < 0.0001). Pairwise differences among cell types were evaluated with Tukey adjustment for multiple comparisons, and odds ratios (ORs) with 95% confidence intervals were reported. For visualization, predicted probabilities and confidence limits were plotted with ggplot2. Each black point represents the model-estimated mean probability for a given neuronal class, error bars denote 95% Wald confidence intervals, and faint gray points indicate observed block rates within individual experiments. This approach provides both a population-level estimate and a visual impression of inter-experiment variability.

### Fish behavioral assays

All animal assays were conducted in accordance with the *Guide for the Care and Use of Laboratory Animals* and were approved by the Institutional Animal Care and Use Committee (IACUC) of the University of Utah (protocol #001629). Goldfish (*Carassius auratus*) of both sexes (weight= 0.85 ± 0.11 g, mean ± SD) were obtained from a local supplier and used for the experiments. An in-house prepared E3 Buffer (4.96 mM NaCl, 0.18 mM KCl, 0.33 mM CaCl₂·2H₂O, 0.4 mM MgCl₂·6H₂O; pH 7.2) served as the vehicle. Groups of three fish each received injections of either 13.5 µL E3 buffer alone (control) or E3 buffer containing the desired concentration of conkazal-M1.

All injections were performed with a 1 mL insulin syringe (31 G). For intraperitoneal injections, fish were positioned ventral side up on a chilled sponge embedded with E3 Buffer under a dissecting microscope, and the solution was delivered into the abdominal cavity. For intramuscular injections, fish were positioned laterally on a hard surface moistened with E3 buffer under a dissecting microscope, and the solution was administered into the lateral muscle below the posterior end of the dorsal fin. Following injection, all fish were maintained under observation for three hours.

## ABBREVIATIONS

CLANS, CLuster ANalysis of Sequences); csPDI, conotoxin-specific PDI; DRG: dorsal root ganglion, DTT, dithiothreitol; ER, endoplasmic reticulum; HEPES, 4-(2-hydroxyethyl)-1-piperazineethanesulfonic acid; hPDI, human PDI; IMAC, immobilized metal affinity chromatography; IPTG, isopropyl ß-D-1-thiogalactopyranoside; KPI: Kazal-type protease inhibitor, LB, lysogeny broth: MSA, multiple sequence alignment; NCBI, National Center for Biotechnology Information; PDB, Protein Data Bank; PDI, protein disulfide isomerase; RMSD, root mean square deviation; SDS-PAGE, sodium dodecyl sulfate-polyacrylamide gel electrophoresis; SRA, sequence read archive; TEVp, tobacco etch virus protease; TPM, transcripts per million; Ub, ubiquitin; Ub-His_10_, Ub containing 10 consecutive histidines.

## ACKNOWLEDGMENTS

We thank Signe A. Sjørup, SBiNLab, University of Copenhagen, for recording mass spectra and Dr. Paula Florez Salcedo, Genentech Inc., for kindly providing illustrations to Figs. 1 and S1.

## AUTHOR CONTRIBUTIONS

Conceptualization, C.M.H., T.L.K., H.S.-H., and L.E.; Data curation, C.M.H., T.L.K., A.R., K.C., M.W., K.T., and L.E.; Formal Analysis, C.M.H., T.L.K., A.R., K.C., M.L.G., and K.T.; Funding acquisition, T.L.K., B.O., H.S.-H., K.T., and L.E.; Investigation, C.M.H., T.L.K., N.L.R., M.L.G., S.E., Z.G.A., M.W., A.R., K.T. and L.E.; Project administration, L.E.; Supervision, B.O., H.S.-H., and L.E.; Visualization, C.M.H., T.L.K., A.R., K.T., and L.E.; writing—original draft preparation, C.M.H., T.L.K., A.R., H.S.-H., K.T. and L.E.; writing—review and editing, all authors. All authors have read and agreed to the published version of the manuscript.

## FOOTNOTES

The atomic coordinates for conkazal-M1 are available with the Protein Data Bank under accession number 9SLR and in the Biological Magnetic Resonance Data Bank under ID number 53313.

## REFERENCES

1. Casewell, N. R., Wuster, W., Vonk, F. J., Harrison, R. A., and Fry, B. G. (2013) Complex cocktails: the evolutionary novelty of venoms Trends Ecol Evol 28, 219–229

2. Lewis, R. J., and Garcia, M. L. (2003) Therapeutic potential of venom peptides Nat Rev Drug Discov 2, 790–802

3. Sachetto, A. T. A., and Mackman, N. (2019) Modulation of the mammalian coagulation system by venoms and other proteins from snakes, arthropods, nematodes and insects Thromb Res 178, 145–154

4. Woodward, S. R., Cruz, L. J., Olivera, B. M., and Hillyard, D. R. (1990) Constant and hypervariable regions in conotoxin propeptides The EMBO journal 9, 1015-1020

5. Sollod, B. L., Wilson, D., Zhaxybayeva, O., Gogarten, J. P., Drinkwater, R., and King, G. F. (2005) Were arachnids the first to use combinatorial peptide libraries? Peptides 26, 131–139

6. Jackson, T. N. W., and Koludarov, I. (2020) How the Toxin got its Toxicity Front Pharmacol 11, 574925

7. Bayrhuber, M., Vijayan, V., Ferber, M., Graf, R., Korukottu, J., Imperial, J., et al. (2005) Conkunitzin-S1 is the first member of a new Kunitz-type neurotoxin family: structural and functional characterization J Biol Chem 280, 23766-23770

8. Finol-Urdaneta, R. K., Remedi, M. S., Raasch, W., Becker, S., Clark, R. B., Struver, N. et al. (2012) Block of Kv1.7 potassium currents increases glucose-stimulated insulin secretion EMBO molecular medicine 4, 424–434

9. Droctove, L., Ciolek, J., Mendre, C., Chorfa, A., Huerta, P., Carvalho, C. et al. (2022) A new Kunitz-type snake toxin family associated with an original mode of interaction with the vasopressin 2 receptor Br J Pharmacol 179, 3470-3481

10. Schweitz, H., Heurteaux, C., Bois, P., Moinier, D., Romey, G., and Lazdunski, M. (1994) Calcicludine, a venom peptide of the Kunitz-type protease inhibitor family, is a potent blocker of high-threshold Ca2+ channels with a high affinity for L-type channels in cerebellar granule neurons Proc Natl Acad Sci USA 91, 878–882

11. Mishra, M. (2020) Evolutionary Aspects of the Structural Convergence and Functional Diversification of Kunitz-Domain Inhibitors J Mol Evol 88, 537–548

12. Yuan, C. H., He, Q. Y., Peng, K., Diao, J. B., Jiang, L. P., Tang, X., et al. (2008) Discovery of a distinct superfamily of Kunitz-type toxin (KTT) from tarantulas PloS one 3, e3414

13. Gladkikh, I., Peigneur, S., Sintsova, O., Lopes Pinheiro-Junior, E., Klimovich, A., Menshov, A., et al. (2020) Kunitz-Type Peptides from the Sea Anemone Heteractis crispa Demonstrate Potassium Channel Blocking and Anti-Inflammatory Activities Biomedicines 8

14. Peigneur, S., Billen, B., Derua, R., Waelkens, E., Debaveye, S., Beress, L. et al. (2011) A bifunctional sea anemone peptide with Kunitz type protease and potassium channel inhibiting properties Biochem Pharmacol 82, 81–90

15. Zupunski, V., and Kordis, D. (2016) Strong and widespread action of site-specific positive selection in the snake venom Kunitz/BPTI protein family Sci Rep 6, 37054

16. Zdenek, C. N., Cardoso, F. C., Robinson, S. D., Mercedes, R. S., Raidjoe, E. R., Hernandez-Vargas, M. J., et al. (2024) Venom exaptation and adaptation during the trophic switch to blood-feeding by kissing bugs iScience 27, 110723

17. Friedrich, T., Kroger, B., Bialojan, S., Lemaire, H. G., Hoffken, H. W., Reuschenbach, P., et al. (1993) A Kazal-type inhibitor with thrombin specificity from Rhodnius prolixus J Biol Chem 268, 16216-16222

18. Kim, B. Y., Lee, K. S., Zou, F. M., Wan, H., Choi, Y. S., Yoon, H. J. et al. (2013) Antimicrobial activity of a honeybee (Apis cerana) venom Kazal-type serine protease inhibitor Toxicon 76, 110–117

19. Liao, Z., Tang, X., Chen, W., Jiang, X., Chen, Z., He, K. et al. (2022) Shrew’s venom quickly causes circulation disorder, analgesia and hypokinesia Cell Mol Life Sci 79, 35

20. Qian, C., Fang, Q., Wang, L., and Ye, G. Y. (2015) Molecular Cloning and Functional Studies of Two Kazal-Type Serine Protease Inhibitors Specifically Expressed by Nasonia vitripennis Venom Apparatus Toxins 7, 2888-2905

21. Wang, Q. W., Zou, W. B., Masson, E., Ferec, C., Liao, Z., and Chen, J. M. (2025) Genetics and clinical implications of SPINK1 in the pancreatitis continuum and pancreatic cancer Hum Genomics 19, 32

22. Brillard-Bourdet, M., Hamdaoui, A., Hajjar, E., Boudier, C., Reuter, N., Ehret-Sabatier, L. et al. (2006) A novel locust (Schistocerca gregaria) serine protease inhibitor with a high affinity for neutrophil elastase Biochem J 400, 467–476

23. Rimphanitchayakit, V., and Tassanakajon, A. (2010) Structure and function of invertebrate Kazal-type serine proteinase inhibitors Dev Comp Immunol 34, 377–386

24. Safavi-Hemami, H., Foged, M. M., and Ellgaard, L. (2018) Evolutionary Adaptations to Cysteine-rich Peptide Folding In Oxidative Folding of Proteins: Basic Principles, Cellular Regulation and Engineering, Feige M, ed. Royal Society of Chemistry, 99–128

25. Koch, T. L., Robinson, S. D., Salcedo, P. F., Chase, K., Biggs, J., Fedosov, A. E., et al. (2024) Prey Shifts Drive Venom Evolution in Cone Snails Mol Biol Evol 41.

26. Robinson, S. D., Safavi-Hemami, H., McIntosh, L. D., Purcell, A. W., Norton, R. S., and Papenfuss, A. T. (2014) Diversity of conotoxin gene superfamilies in the venomous snail, Conus victoriae PloS one 9, e87648

27. Hackney, C. M., Florez Salcedo, P., Müller, E., Koch, T. L., Kjelgaard, L. D., Watkins, M., et al. (2023) A previously unrecognized superfamily of macro-conotoxins includes an inhibitor of the sensory neuron calcium channel Cav2.3 PLoS Biol 21, e3002217

28. Yang, Z. (1997) PAML: a program package for phylogenetic analysis by maximum likelihood Comput Appl Biosci 13, 555–556

29. Bertelsen, A. B., Hackney, C. M., Bayer, C. N., Kjelgaard, L. D., Rennig, M., Christensen, B. et al. (2021) DisCoTune: versatile auxiliary plasmids for the production of disulphide-containing proteins and peptides in the E. coli T7 system Microb Biotechnol 10.1111/1751-7915.13895

30. Gaciarz, A., Khatri, N. K., Velez-Suberbie, M. L., Saaranen, M. J., Uchida, Y., Keshavarz-Moore, E. et al. (2017) Efficient soluble expression of disulfide bonded proteins in the cytoplasm of Escherichia coli in fed-batch fermentations on chemically defined minimal media Microb Cell Fact 16, 108

31. Nielsen, L. D., Foged, M. M., Albert, A., Bertelsen, A. B., Soltoft, C. L., Robinson, S. D. et al. (2019) The three-dimensional structure of an H-superfamily conotoxin reveals a granulin fold arising from a common ICK cysteine framework J Biol Chem 294, 8745-8759

32. Safavi-Hemami, H., Li, Q., Jackson, R. L., Song, A. S., Boomsma, W., Bandyopadhyay, P. K. et al. (2016) Rapid expansion of the protein disulfide isomerase gene family facilitates the folding of venom peptides Proc Natl Acad Sci USA 113, 3227-3232

33. Rogov, V. V., Rozenknop, A., Rogova, N. Y., Lohr, F., Tikole, S., Jaravine, V. et al. (2012) A universal expression tag for structural and functional studies of proteins Chembiochem 13, 959–963

34. Buczek, O., Bulaj, G., and Olivera, B. M. (2005) Conotoxins and the posttranslational modification of secreted gene products Cell Mol Life Sci 62, 3067-3079

35. Fischer, W. H., and Spiess, J. (1987) dentification of a mammalian glutaminyl cyclase converting glutaminyl into pyroglutamyl peptides Proc Natl Acad Sci USA 84, 3628-3632

36. van Kempen, M., Kim, S. S., Tumescheit, C., Mirdita, M., Lee, J., Gilchrist, C. L. M. et al. (2024) Fast and accurate protein structure search with Foldseek Nat Biotechnol 42, 243–246

37. Krissinel, E., and Henrick, K. (2004) Secondary-structure matching (SSM), a new tool for fast protein structure alignment in three dimensions Acta Crystallogr D Biol Crystallogr 60, 2256-2268

38. Derache, C., Epinette, C., Roussel, A., Gabant, G., Cadene, M., Korkmaz, B. et al. (2012) Crystal structure of greglin, a novel non-classical Kazal inhibitor, in complex with subtilisin FEBS J 279, 4466-4478

39. Perrakis, A., and Sixma, T. K. (2021) AI revolutions in biology: The joys and perils of AlphaFold EMBO Rep 22, e54046

40. Shenoy, R. T., Thangamani, S., Velazquez-Campoy, A., Ho, B., Ding, J. L., and Sivaraman, J. (2011) Structural basis for dual-inhibition mechanism of a non-classical Kazal-type serine protease inhibitor from horseshoe crab in complex with subtilisin PloS one 6, e18838

41. Hordijk, P. L., de Jong-Brink, M., ter Maat, A., Pieneman, A. W., Lodder, J. C., and Kits, K. S. (1992) The neuropeptide schistosomin and haemolymph from parasitized snails induce similar changes in excitability in neuroendocrine cells controlling reproduction and growth in a freshwater snail Neurosci Lett 136, 193–197

42. Frickey, T., and Lupas, A. (2004) CLANS: a Java application for visualizing protein families based on pairwise similarity Bioinformatics 20, 3702-3704

43. Tian, M., Benedetti, B., and Kamoun, S. (2005) A Second Kazal-like protease inhibitor from Phytophthora infestans inhibits and interacts with the apoplastic pathogenesis-related protease P69B of tomato Plant Physiol 138, 1785-1793

44. Tian, M., and Kamoun, S. (2005) A two disulfide bridge Kazal domain from Phytophthora exhibits stable inhibitory activity against serine proteases of the subtilisin family BMC Biochem 6, 15

45. Li, X. C., Wang, X. W., Wang, Z. H., Zhao, X. F., and Wang, J. X. (2009) A three-domain Kazal-type serine proteinase inhibitor exhibiting domain inhibitory and bacteriostatic activities from freshwater crayfish Procambarus clarkii Dev Comp Immunol 33, 1229-1238

46. Esmaeilishirazifard, E., Usher, L., Trim, C., Denise, H., Sangal, V., Tyson, G. H. et al. (2022) Bacterial Adaptation to Venom in Snakes and Arachnida Microbiol Spectr 10, e0240821

47. De Jong-Brink, M. (1995) How schistosomes profit from the stress responses they elicit in their hosts Adv Parasitol 35, 177–256

48. Zhang, S. M., Nian, H., Wang, B., Loker, E. S., and Adema, C. M. (2009) Schistosomin from the snail Biomphalaria glabrata: expression studies suggest no involvement in trematode-mediated castration Mol Biochem Parasitol 165, 79–86

49. Force, A., Lynch, M., Pickett, F. B., Amores, A., Yan, Y. L., and Postlethwait, J. (1999) Preservation of duplicate genes by complementary, degenerative mutations Genetics 151, 1531-1545

50. van de Locht, A., Lamba, D., Bauer, M., Huber, R., Friedrich, T., Kroger, B. et al. (1995) Two heads are better than one: crystal structure of the insect derived double domain Kazal inhibitor rhodniin in complex with thrombin The EMBO journal 14, 5149-5157

51. Maxwell, M., Undheim, E. A. B., and Mobli, M. (2018) Secreted Cysteine-Rich Repeat Proteins “SCREPs”: A Novel Multi-Domain Architecture Front Pharmacol 9, 1333

52. Liu, J., Maxwell, M., Cuddihy, T., Crawford, T., Bassetti, M., Hyde, C., et al. (2023) ScrepYard: An online resource for disulfide-stabilized tandem repeat peptides Protein Sci 32, e4566

53. Katoh, K., Misawa, K., Kuma, K., and Miyata, T. (2002) MAFFT: a novel method for rapid multiple sequence alignment based on fast Fourier transform Nucleic Acids Res 30, 3059-3066

54. Katoh, K., and Standley, D. M. (2013) MAFFT multiple sequence alignment software version 7: improvements in performance and usability Mol Biol Evol 30, 772–780

55. Waterhouse, A. M., Procter, J. B., Martin, D. M., Clamp, M., and Barton, G. J. (2009) Jalview Version 2--a multiple sequence alignment editor and analysis workbench Bioinformatics 25, 1189-1191

56. Slater, G. S., and Birney, E. (2005) Automated generation of heuristics for biological sequence comparison BMC Bioinformatics 6, 31

57. Minh, B. Q., Schmidt, H. A., Chernomor, O., Schrempf, D., Woodhams, M. D., von Haeseler, A. et al. (2020) IQ-TREE 2: New Models and Efficient Methods for Phylogenetic Inference in the Genomic Era Mol Biol Evol 37, 1530-1534

58. Hoang, D. T., Chernomor, O., von Haeseler, A., Minh, B. Q., and Vinh, L. S. (2018) UFBoot2: Improving the Ultrafast Bootstrap Approximation Mol Biol Evol 35, 518–522

59. Rambaut, A. (2018) FigTree (Version 1.4.4) Institute of Evolutionary Biology, University of Edinburgh.

60. Khilji, M. S., Hackney, C. M., Koch, T. L., Hone, A. J., Rogalski, A., Watkins, M., et al. (2025) Structural similarities reveal an expansive conotoxin family with a two-finger toxin fold Protein Sci 34, e70333

61. Wilkins, M. R., Gasteiger, E., Bairoch, A., Sanchez, J. C., Williams, K. L., Appel, R. D. et al. (1999) Protein identification and analysis tools in the ExPASy server Methods Mol Biol 112, 531–552

62. Kazimierczuk, K., and Orekhov, V. Y. (2011) Accelerated NMR spectroscopy by using compressed sensing Angew Chem Int Ed Engl 50, 5556-5559

63. Delaglio, F., Grzesiek, S., Vuister, G. W., Zhu, G., Pfeifer, J., and Bax, A. (1995) NMRPipe: a multidimensional spectral processing system based on UNIX pipes J Biomol NMR 6, 277–293

64. Skinner, S. P., Fogh, R. H., Boucher, W., Ragan, T. J., Mureddu, L. G., and Vuister, G. W. (2016) CcpNmr AnalysisAssign: a flexible platform for integrated NMR analysis J Biomol NMR 66, 111–124

65. Güntert, P., and Buchner, L. (2015) Combined automated NOE assignment and structure calculation with CYANA J Biomol NMR 62, 453–471

66. Schwieters, C. D., Kuszewski, J. J., Tjandra, N., and Clore, G. M. (2003) The Xplor-NIH NMR molecular structure determination package J Magn Reson 160, 65–73

67. Bermejo, G. A., Clore, G. M., and Schwieters, C. D. (2012) Smooth statistical torsion angle potential derived from a large conformational database via adaptive kernel density estimation improves the quality of NMR protein structures Protein Sci 21, 1824-1836

68. Tian, Y., Schwieters, C. D., Opella, S. J., and Marassi, F. M. (2017) High quality NMR structures: a new force field with implicit water and membrane solvation for Xplor-NIH J Biomol NMR 67, 35–49

69. Laskowski, R. A., Rullmannn, J. A., MacArthur, M. W., Kaptein, R., and Thornton, J. M. (1996) AQUA and PROCHECK-NMR: programs for checking the quality of protein structures solved by NMR J Biomol NMR 8, 477–486

70. Olp, M. D., Kalous, K. S., and Smith, B. C. (2020) ICEKAT: an interactive online tool for calculating initial rates from continuous enzyme kinetic traces BMC Bioinformatics 21, 186

71. Teichert, R. W., Smith, N. J., Raghuraman, S., Yoshikami, D., Light, A. R., and Olivera, B. M. (2012) Functional profiling of neurons through cellular neuropharmacology Proc Natl Acad Sci USA 109, 1388-1395

