## Supplementary figures for "Conkazal-M1 from the MKAVA family of conotoxins – a dual-function protease inhibitor and neuroactive peptide"

***Correspondence:**

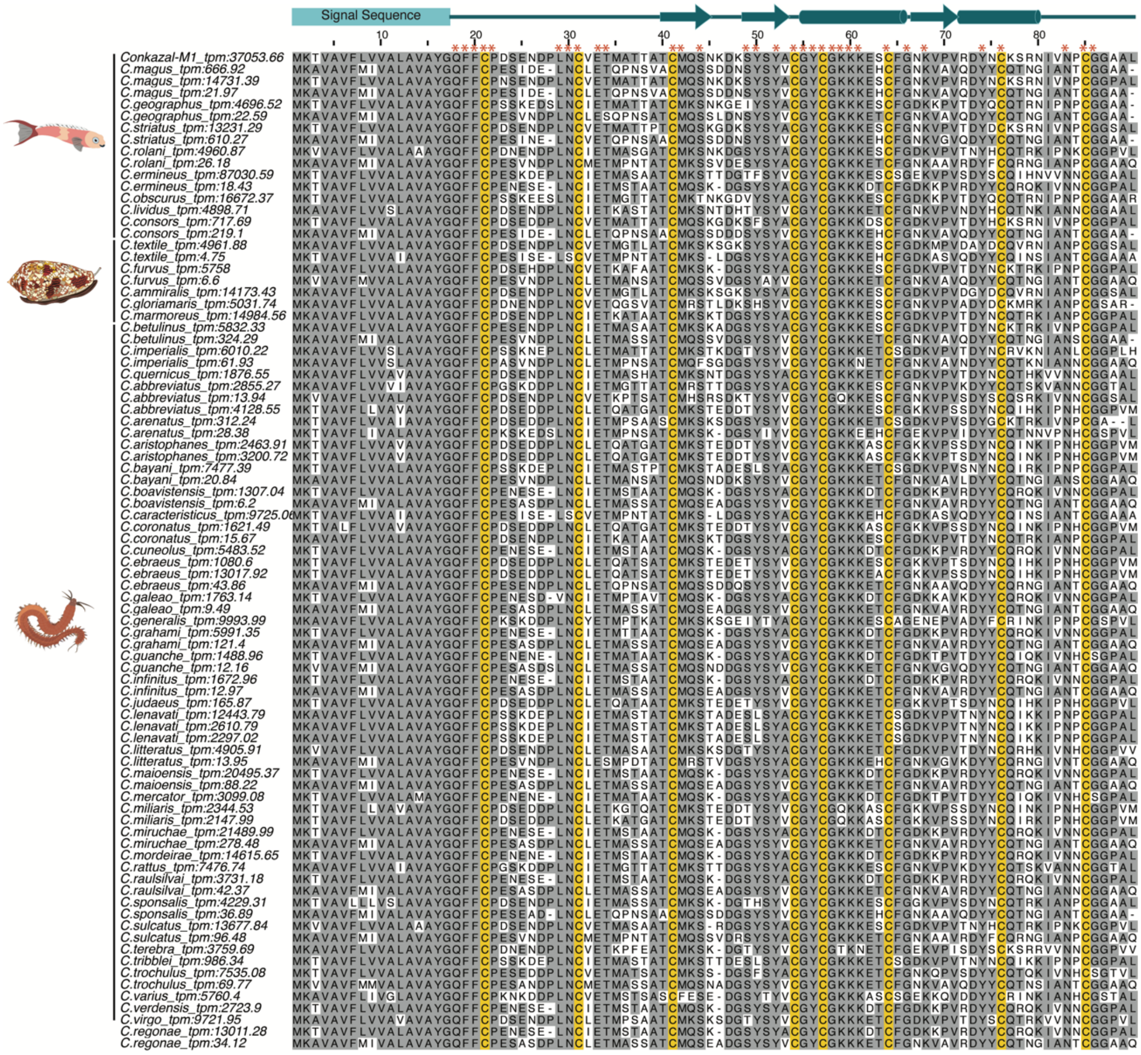


**Figure S1. Multiple sequence alignment of MKAVA superfamily conotoxins**

The light teal N-terminal rectangle indicates the position of the predicted signal sequences, while the dark teal arrows and cylinders above the sequences show the positions of β-strands and α-helices in the mature toxin, as determined by NMR spectroscopy in the mature toxin sequence. Cysteine residues are highlighted in yellow. Residues highlighted in grey share at least 40% identity at the given position in the alignment, and red asterisks denote residues in the mature region conserved in ≥95% of sequences across the entire family. The prey preference for each species (fish, other molluscs, or worms) is indicated by the icons on the left, kindly provided by Dr. Paula Florez Salcedo.

***Conus marmoreus***
MKAVAVFLVVALAVAYGQFFCPDSENDPLNCIETK**ATAATCMKSKTDGSYSYACGYCGKKKESCFGDKVPVTDYNCQTRKIANPCGGPAL**

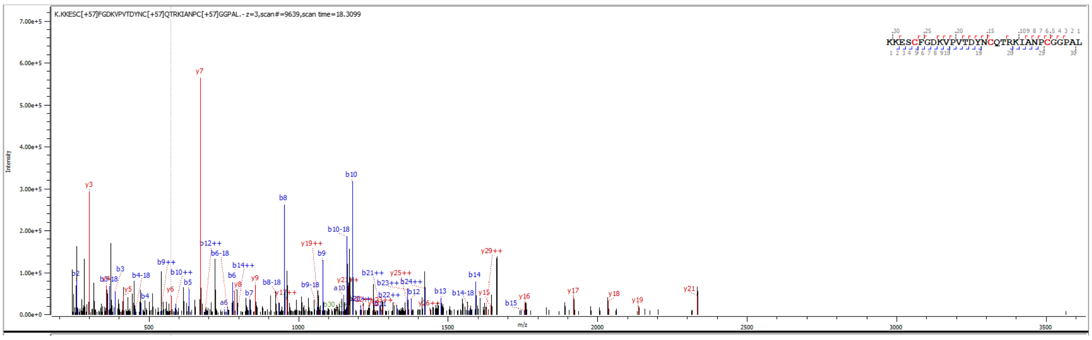


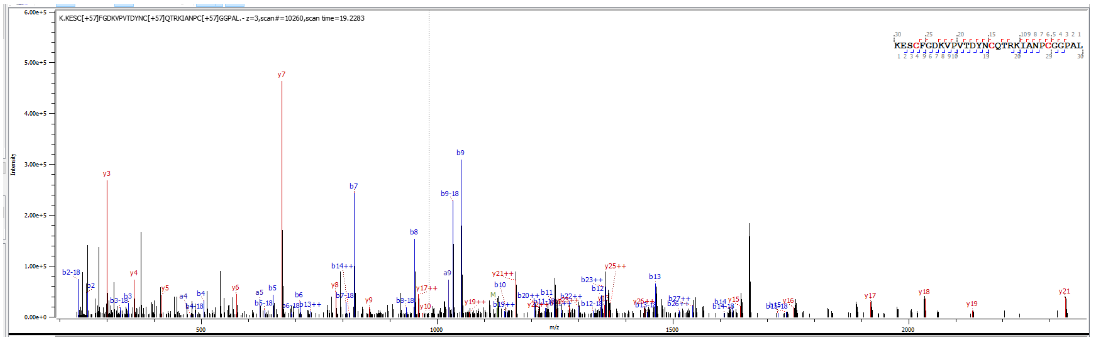


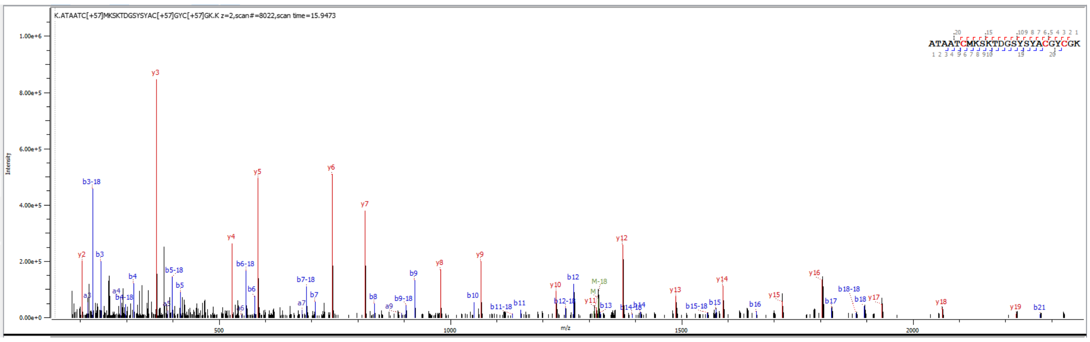


***Conus geographus***
MKTVAVFLVVALAVAYGQFFCPSSKEDSLNCIETMATTATCMKSNKGEIYSYACGYCGKKKESCFGDK**KPVTDYQCQTR**NIPNPCGGAA

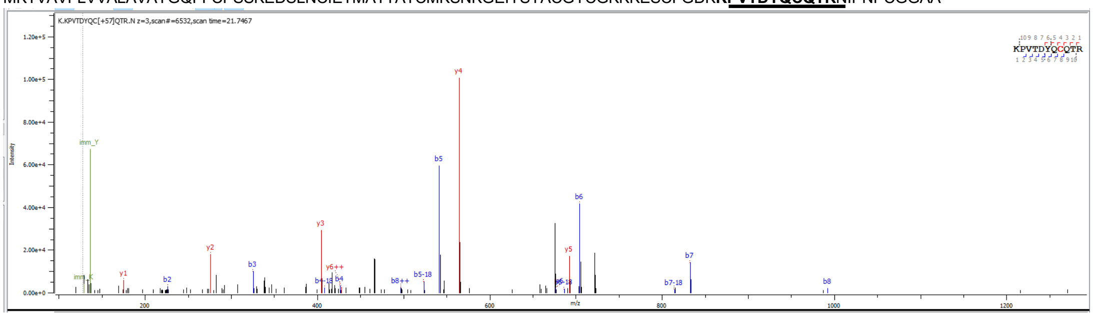


### Figure S2. Proteomics-based identification of MKAVA peptides in venom

Tryptic peptides identified from the venom of *Conus geographus* and *Conus marmoreus* using Byonic software. The depicted spectra were manually verified.

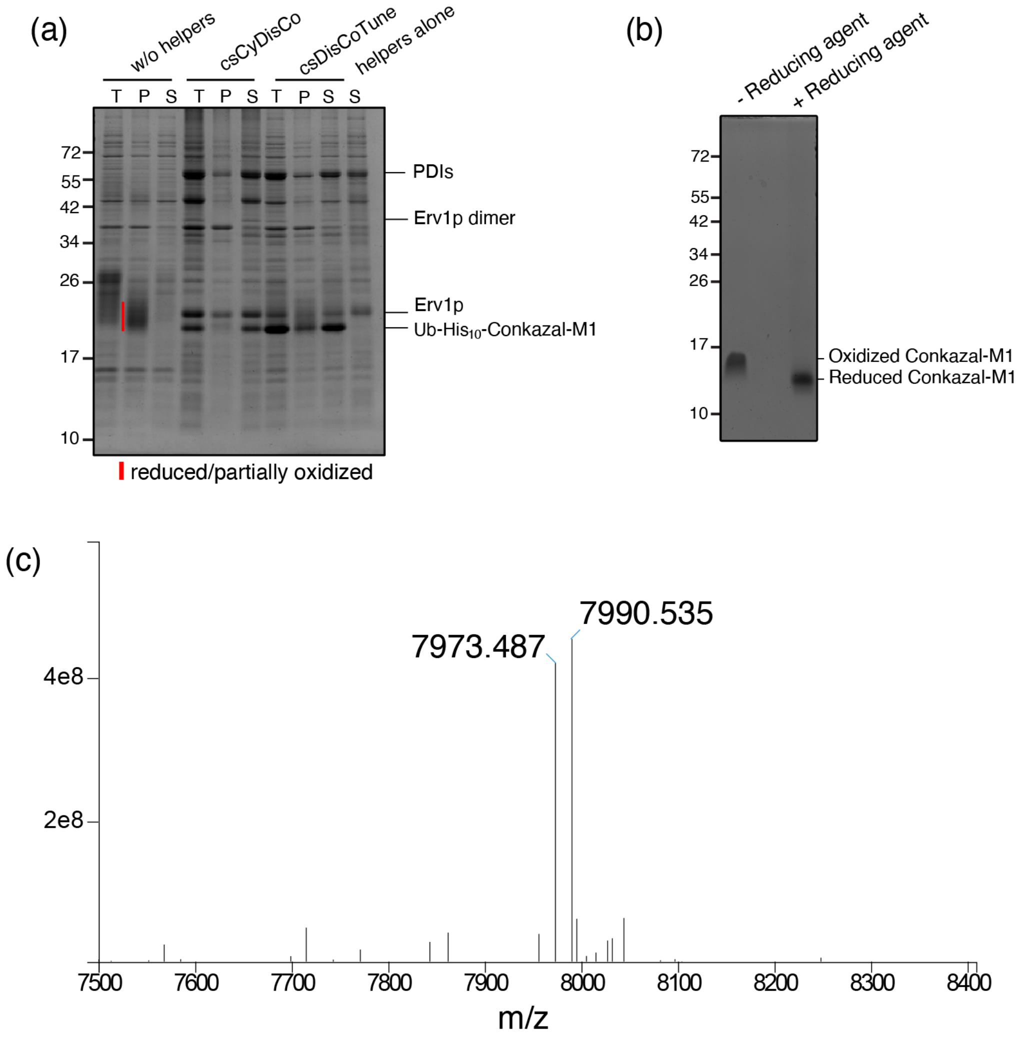


**Figure S3. Conkazal-M1 is efficiently produced in *E. coli* using the csDisCoTune system.**

**(a)** SDS-PAGE analysis of Ub-$His_{10}$-Conkazal-M1 expressed in *E. coli* BL21(DE3) cells without (w/o helpers), with the csCyDisCo system (csCyDisCo) or with the csDisCoTune system (csDisCoTune). Samples from the total cell extract (T), the soluble fraction (S), and the insoluble fraction solubilized in urea (P) were analyzed by SDS-PAGE. Protein levels have been normalized and are directly comparable across lanes. **(b)** 15% SDS-PAGE gel analysis of purified Conkazal-M1 in the reduced (40 mM DTT; +reducing agent) and oxidized (-reducing agent) state. Molecular mass markers (in kDa) are indicated on the left. The protein bands were visualized with Coomassie Brilliant Blue. **(c)** Q-TOF mass spectrum for purified conkazal-M1 showing a dominant peak at a monoisotopic mass of 7990.54 Da. The theoretical monoisotopic mass for oxidized conkazal-M1 is 7990.55 Da. The secondary peak exhibits a monoisotopic mass of

7973.49 Da consistent with a post-translational modification of the N-terminal Gln to a pyroglutamate.

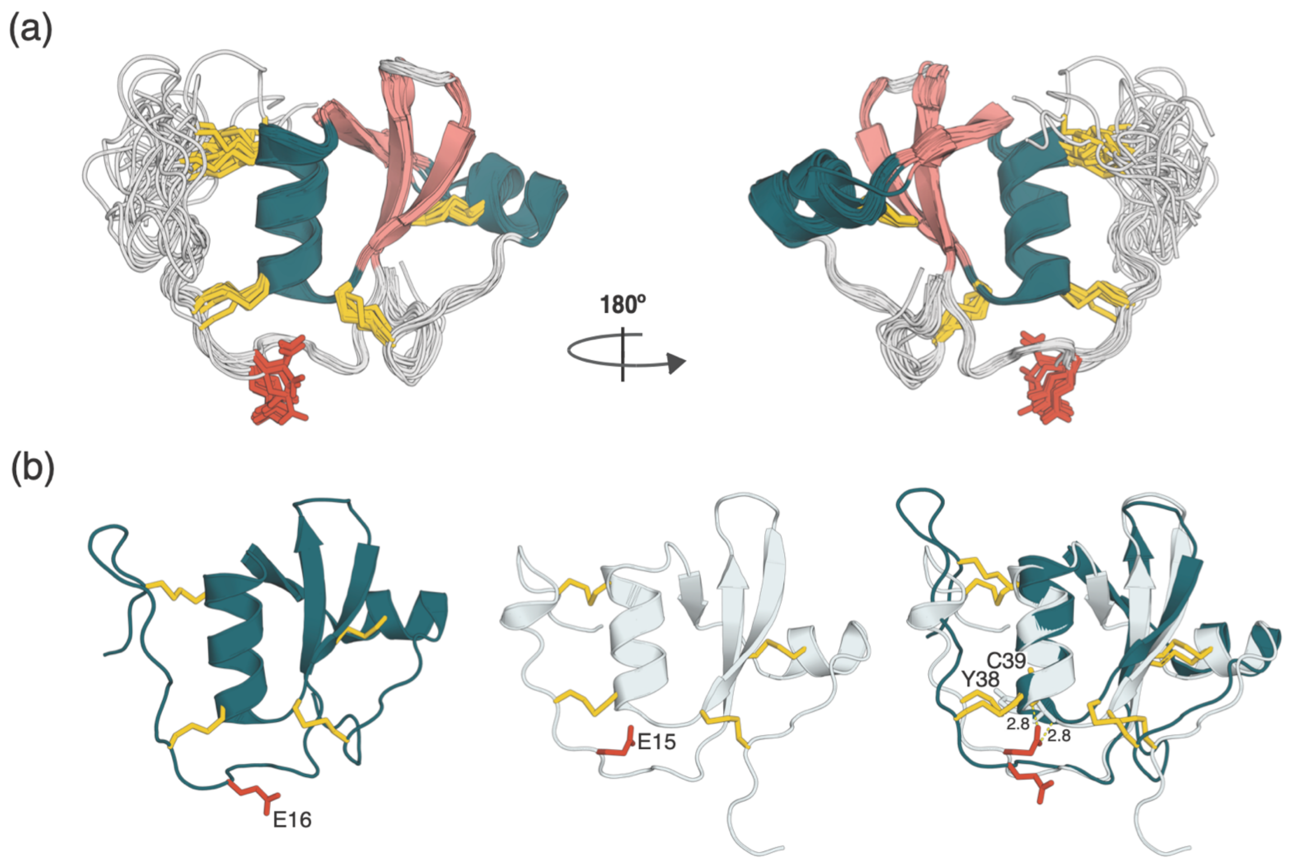


**Figure S4. The structure of conkazal-M1 is well-ordered from residue 10**

(**a**) Cartoon representation of the NMR structure of conkazal-M1 (PDB: 9SLR) shown from two different angles. The 20 lowest energy structures of conkazal-M1 are superimposed. Cysteine residues are displayed in yellow, the $\alpha$-helix in teal, and $\beta$-strands in pink. The conserved glutamate at position 16 is shown as red sticks. (**b**) Cartoon representation of a single conformer of the conkazal-M1 NMR structure (left), cartoon representation of the Alphafold3 prediction of Conkazal-S1 from *C. striatus* (middle), and superposition of conkazal-M1 and Conkazal-S1 (right). Disulfide bridges are colored yellow and the amino acid residue at the P1 position is shown in red. The hydrogen bonds predicted to stabilize the Glu15 of Conkazal-S1 are shown as yellow, dotted lines (with distances indicated), while the two amino acids (Tyr38 and Cys39) predicted to coordinate the H-bonds are shown as sticks.

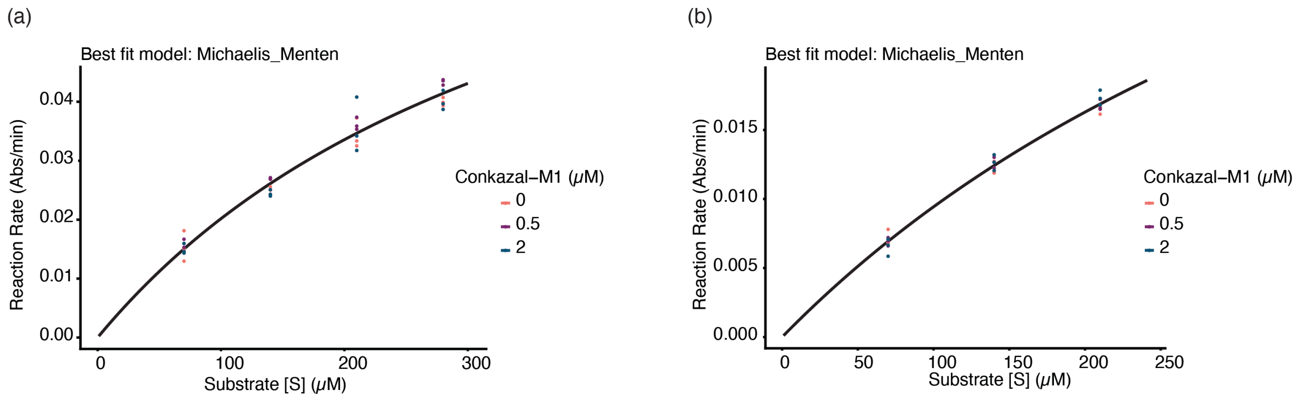


**Figure S5. Conkazal-M1 does not inhibit the activity of chymotrypsin or GluC.**

Kinetic analysis of (a) GluC and (b) chymotrypsin in the presence of increasing concentrations of conkazal-M1. Reaction rates were measured across a range of substrate concentrations and were normalized to enzyme mass. The plots display experimental data points (colored according to inhibitor concentration: 0 mM (orange), 0.5 mM (purple), 2 mM (dark teal)) overlaid with lines showing the best-fit model (Michaelis-Menten, i.e., no inhibition). The reaction rate is shown in absorption (Abs) per minute.

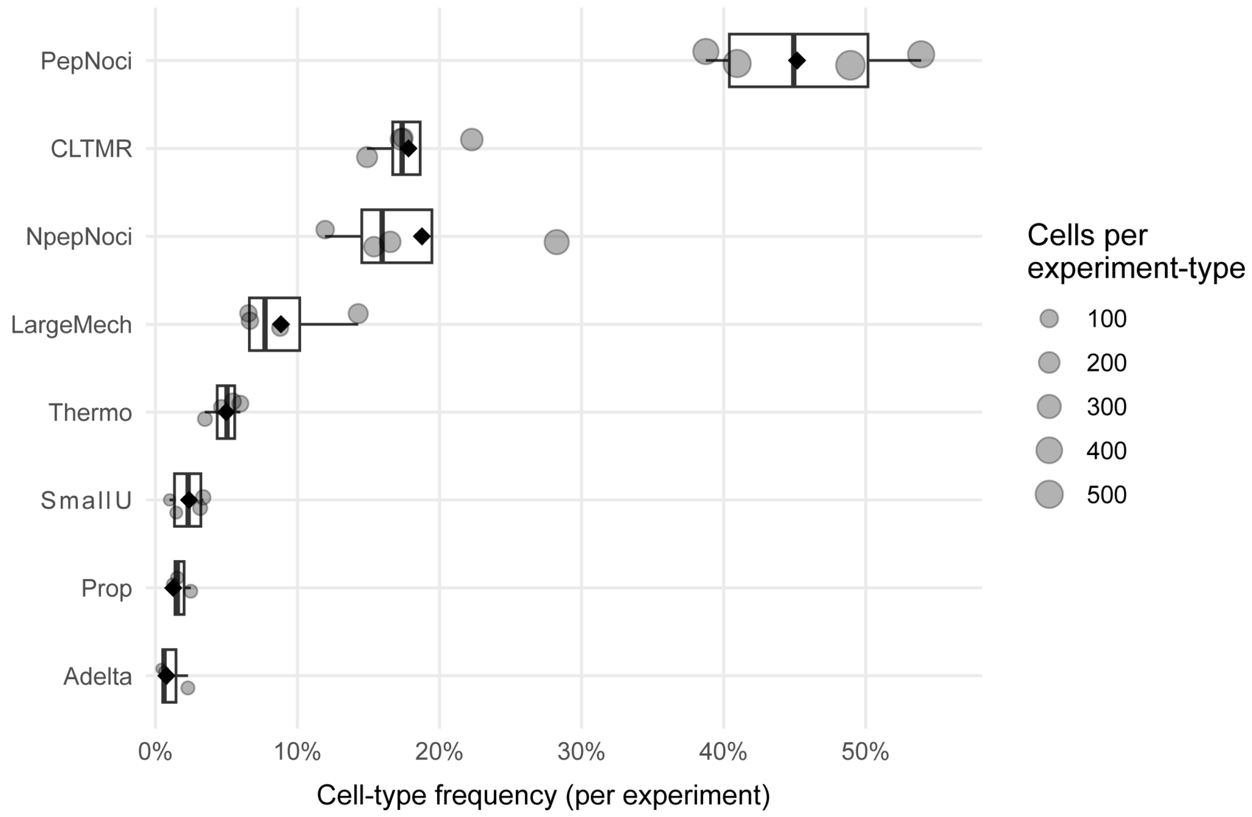


**Figure S6. Cell-type composition across experiments**

For each experiment, the proportion of cells belonging to each neuronal class was calculated as the number of cells of that type divided by the total number of cells recorded in that experiment. Experiments in which a given cell type was not observed were included with a frequency of zero to accurately represent variability in detection and true absences. Boxplots show the distribution of per-experiment frequencies for each neuronal class, and jittered points indicate individual experiment values (point size ∝ number of cells of that type in that experiment). Black diamonds denote the pooled frequency of each cell type computed across all experiments. This analysis demonstrates the stability and variability of sensory neuron class representation across biological replicates.

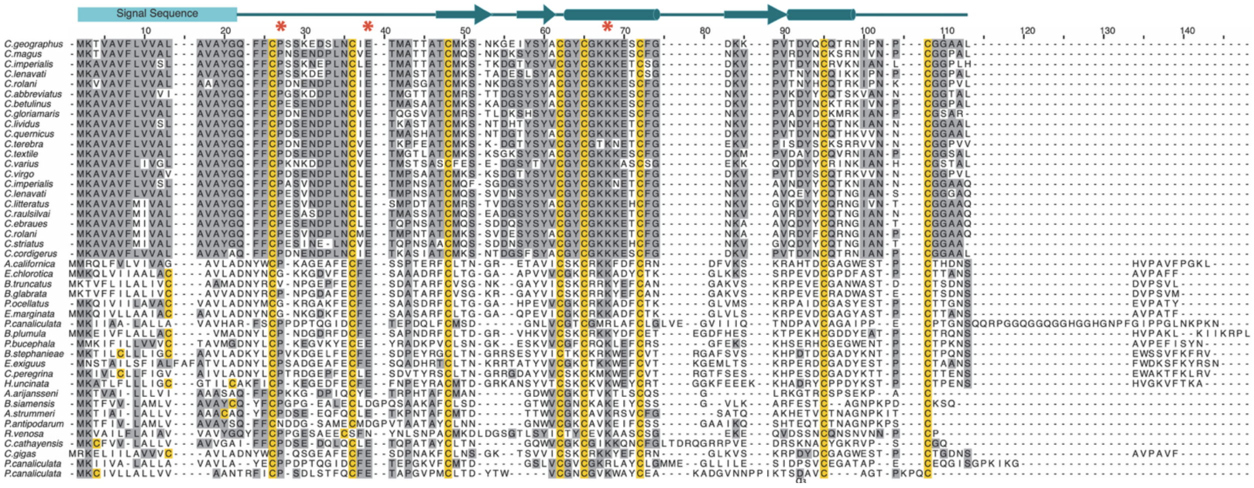


**Figure S7. Multiple sequence alignments of MKAVA and schistosomin sequences**

The position of the predicted signal sequence and the regular secondary structure elements in Conkazal-M1 are indicated above the alignment, as in Figure S1. Residues highlighted in grey indicate at least 30% identity at the given position in the alignment, except for the cysteines, which are highlighted in yellow regardless of percent identity. Sequences for alignment were selected from a larger dataset by excluding those with >90% similarity, retaining only sequences with < 90% similarity for display. Red asterisks denote amino acid residues within the mature protein region exhibiting over 88% identity across all MKAVA and schistosomin sequences analyzed.

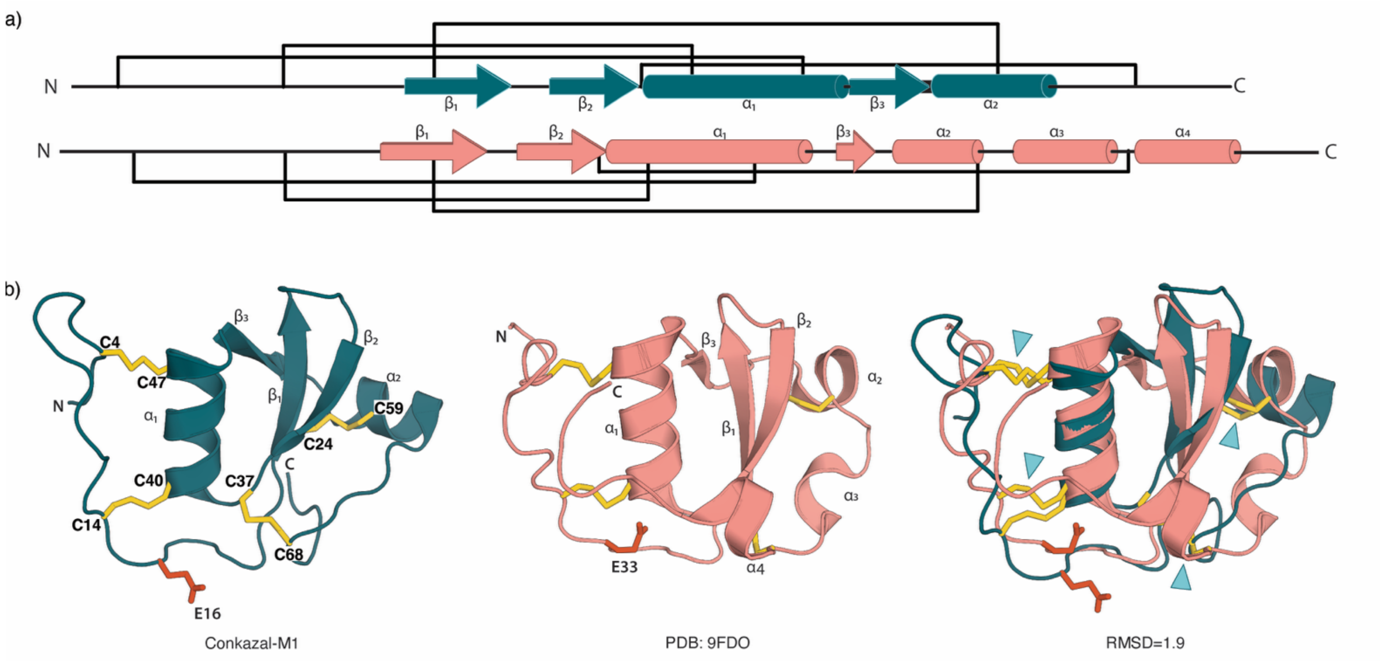


**Figure S8.** Conkazal-M1 harbors a similar fold to schistosomin. (**a**). Graphic representations of Conkazal-M1 (teal) and schistosomin from *Biomphalaria glabrata* (pink). α-helices and β-strands are represented by cylinders and block arrows, respectively, and disulfide bridges are represented as black brackets. (**b**). Cartoon representation of the conkazal-M1 NMR structure (PDB: 9SLR) (left), cartoon representation of the *B. glabrata* schistosomin structure solved by X-ray crystallography (PDB: 9FDO) (middle), and superposition of conkazal-M1 and schistosomin (right). Coloration as described in Panel (a). Disulfide bridges are colored yellow, and the amino acid residue at the P1 position is shown in red. The three conserved disulfide bridges are labeled with blue arrowheads (right). The RMSD value is derived from 38 α-carbons and determined using the PyMOL "super" command.

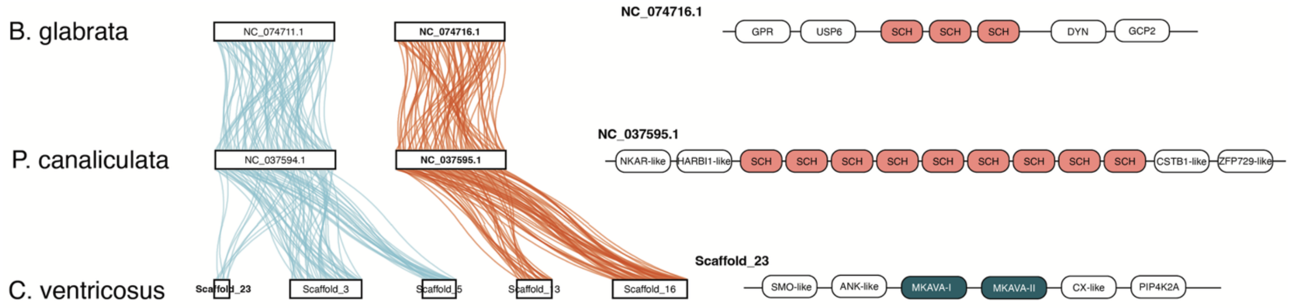


**Figure S9. Synteny analyses indicate that conkazals and schistosomins belong to distinct linkage groups.**

(Left) Ribbon diagrams of orthologous proteins from *C. ventricosus*, *P. canaliculata*, and *B. glabrata* reveal homologous chromosomes across the three gastropods. The conkazal genes of *C. ventricosus* are positioned on pseudochromosomes that are not homologous to those containing schistosomin genes in *P. canaliculata* and *B. glabrata*. (Right) Both conkazals and schistosomins occur as tandem duplicated paralogs on their respective chromosomes. However, they lack conserved microsynteny with neighboring genes.

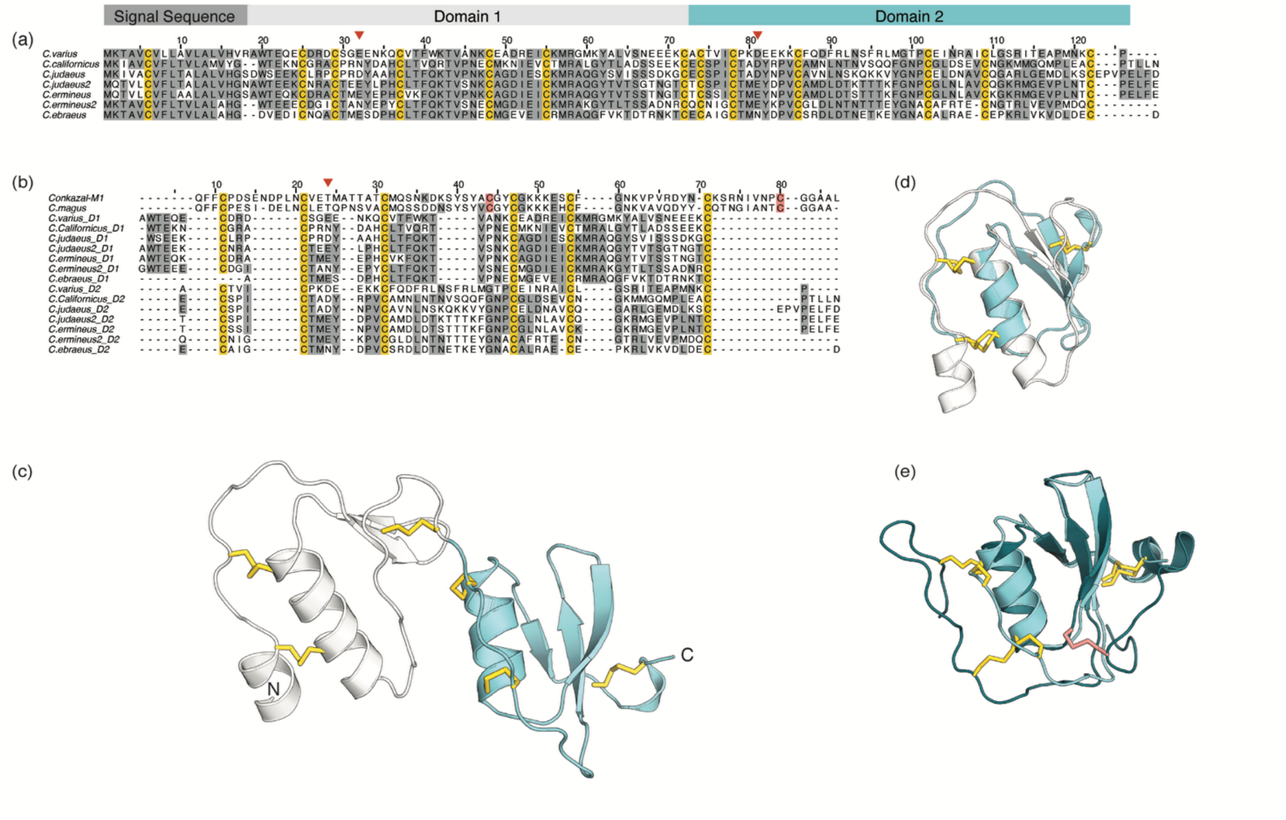


**Figure S10. Sequences of the MKTAV family of conotoxins represent two-domain kazal-type proteins**

**(a)** Alignment of MKTAV superfamily sequences from different cone snail species. Residues highlighted in gray indicate at least ≥40% identity at the given position in the alignment. Red arrowheads above the alignment designate the amino acid residues in the proposed non-canonical P1 positions. **(b)** Alignment of conkazal-M1 and *C. magus* conkazals with MKTAV sequences split into the individual domains (D1 and D2). **(c)** Alphafold3 prediction of MKTAV from *Conus varius*, with domain 1 (D1) in white and domain 2 (D2) in light blue. **(d)** Structural overlay of MKTAV D1 (white) and D2 (light blue). **(e)** Overlay of MKTAV D2 (light blue) with conkazal-M1 (dark teal). Disulfide bonds are shown in yellow, except the additional disulfide in conkazal-M1, which is depicted in pink.

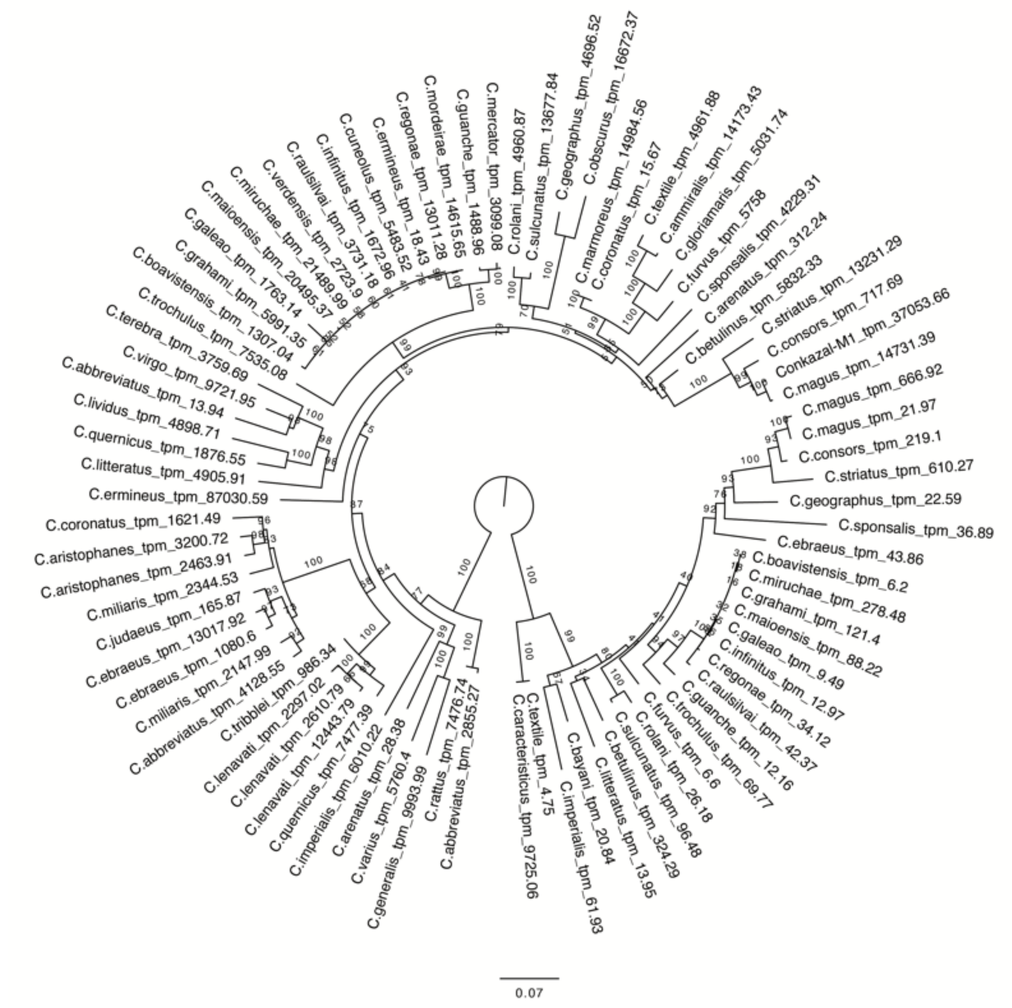


**Figure S11. Phylogenetic reconstruction from Fig. 1(c) with all bootstrap values present.** Maximum-likelihood was performed using IQ-Tree's UF bootstrap with 1000 replicates, with best fit model TMP3 + G4 selected by the Bayesian Information Criterion. The tree was rooted to separate the two paralogous genes. The scale bar represents genetic distance.
