## Supplementary material for "Conkazal-M1 from the MKAVA family of conotoxins – a dual-function protease inhibitor and neuroactive peptide": Suppl. File 1

**Supplementary File 1.** All full-length conkazal superfamily sequences used for the multiple sequence alignment in Fig. S1.

>Conkazal-M1_tpm:37053.66

MKTVAVFLVVALAVAYGQFFCPDSENDPLNCVETMATTATCMQSNKDKSYSYACGYCGKKKESCFGNKVPVRDYNCKSRNIVNPCGGAAL

>C.magus_tpm:666.92

MKAVAVFMIVALAVAYGQFFCPESIDELNCLETQPNSVACMQSSDDNSYSYVCGYCGKKKEHCFGNKVAVQDYYCQTNGIANTCGGAA

>C.magus_tpm:14731.39

MKTVAVFLVVALAVAYGQFFCPNSENDPLNCVETMATTATCMQSNKDKSYSYACGYCGKKKESCFGNKVPVRDYNCKSRNIVNPCGGPAL

>C.magus_tpm:21.97

MKAVAVFMIVALAVAYGQFFCPESIDELNCLETQPNSVACMQSSDDNSYSYVCGYCGKKKEHCFGNKVAVQDYYCQTNGIANTCGGAA

>C.geographus_tpm:4696.52

MKTVAVFLVVALAVAYGQFFCPSSKEDSLNCIETMATTATCMKSNKGEIYSYACGYCGKKKESCFGDKKPVTDYQCQTRNIPNPCGGAA

>C.geographus_tpm:22.59

MKAVAVFMIVALAVAYGQFFCPESVNDPLNCLESQPNSATCMQSSLDNSYSYVCGYCGKKKETCFGNKVAVQDYYCQTNGIANTCGGAA

>C.striatus_tpm:13231.29

MKTVAVFLVVALAVAYGQFFCPDSENDPLNCVETMATTPTCMQSKGDKSYSYACGYCGKKKESCFGNKVPVTDYDCKSRNIVNPCGGSAL

>C.striatus_tpm:610.27

MKAVAVFMIVALAVAYGQFFCPESINELNCVETQPNSAACMQSSDDNSYSYVCGYCGKKKEHCFGNKVGVQDYYCQTNGIANTCGGAA

>C.rolani_tpm:4960.87

MKVVAVFLVVALAAAYGQFFCPDNENDPLNCIETMASGATCMKSNKDGSYSYACGYCGKKKESCFGDKVPVTNYHCQTRKIPNKCGGPVL

>C.rolani_tpm:26.18

MKAVAVFMIVALAVAYGQFFCPESVNDPLNCMETMPNTATCMKSSVDESYSYACGYCGKKKETCFGNKAAVRDYFCQRNGIANPCGGAAQ

>C.textile_tpm:4961.88

MKAVAVFLVVALAVAYGQFFCPDSENDPLNCVETMGTLATCMKSKSGKSYSYACGYCGKKKESCFGDKMPVDAYDCQVRNIANPCGGSAL

>C.textile_tpm:4.75

MKTVAVFLVVAIAVAYGQFFCPESISELSCVETMPNTATCMKSLDGSYSYACGYCGKKKEHCFGDKASVQDYYCQINSIANTCGGAAA

>C.furvus_tpm:5758

MKAVAVFLVVALAVAYGQFFCPDSEHDPLNCVETKAFAATCMKSKDGSYSYACGYCGKKKESCFGDKVPVTDYNCKTRKIPNPCGGPAL

>C.furvus_tpm:6.6

MKVVAVFMVVALAVAYGQFFCPESVNDPLNCLETMANSATCMQSSVDGSYAYVCGYCGKKKETCFGNKVAVQDYYCQTNGIANTCGGAAQ

>C.ammiralis_tpm:14173.43

MKAVAVFLVVALAVAYGQFFCPDSENDPLNCVETMGTLATCMKSKSGKSYSYACGYCGKKKESCFGDKVPVDGYDCQVRNIANPCGGSAL

>C.gloriamaris_tpm:5031.74

MKAVAVFLVVALAVAYGQFFCPDNENDPLNCVETQGSVATCMRSTLDKSHSYVCGYCGKKKESCFGNKVPVADYDCKMRKIANPCGSAR

>C.marmoreus_tpm:14984.56

MKAVAVFLVVALAVAYGQFFCPDSENDPLNCIETKATAATCMKSKTDGSYSYACGYCGKKKESCFGDKVPVTDYNCQTRKIANPCGGPAL

>C.betulinus_tpm:5832.33

MKAVAVFLVVALAVAYGQFFCPESENDPLNCIETMASAATCMKSKADGSYSYACGYCGKKKESCFGDKVPVTDYNCKTRKIVNPCGGPAL

>C.betulinus_tpm:324.29

MKAVAVFMIVALAVAYGQFFCPESVNDPLNCLETMASSATCMQSSVDGSYSYVCGYCGKKKETCFGNKVAVQDYYCQTNGIANSCGGAA

>C.imperialis_tpm:6010.22

MKAVAVFLVVSLAVAYGQFFCPSSKNEPLNCLETMATTATCMKSTKDGTYSYVCGYCGKKKETCSGDKVPVTDYNCRVKNIANLCGGPLH

>C.imperialis_tpm:61.93

MKAVAVFLVVSLAVAYGQFFCPASVNDPLNCLETMPNSATCMQFSGDGSYSYVCGYCGKKNETCFGNKVAVNDYYCQTKNIANNCGGAAQ

>C.quernicus_tpm:1876.55

MKAVAVFLVVAVAVAYGQFFCPDSENDPLNCIETMASHATCMKSNTDGSYSYACGYCGKKKETCFGDKVPVTDYNCQTHKVVNNCGGAAL

>C.abbreviatus_tpm:2855.27

MKAVAVFLVVVIAVAYGQFFCPGSKDDPLNCIETMGTTATCMRSTTDGSYSYACGYCGKKKESCFGNKVPVKDYYCQTSKVANNCGGTAL

>C.abbreviatus_tpm:13.94

MKVVAVFLVVALAVAYGQFFCPDSENDPLNCVETKPTSATCMHSRSDKTYSYVCGYCGQKKESCFGNKVPVSDYSCQSRKIVNNCGGSAL

>C.abbreviatus_tpm:4128.55

MKTVAVFLLVAVAVAYGQFFCPDSEDDPLNCLETQATGATCMKSTEDDTYSYVCGYCGKKKESCFGKKVPSSDYNCQIHKIPNHCGGPVM

>C.arenatus_tpm:312.24

MKTVAVFLVVAIAVAYGQFFCPDSEDDPLNCIETMPSAASCMKSKSDGSYSYVCGYCGKKKETCSGDKVPVSDYGCKTRKIVNPCGAL

>C.arenatus_tpm:28.38

MKTVAVFLIVALAVAYGQFFCPKSKEDSLNCIETMPNSATCMKSKDGSYIYVCGYCGKKEEHCFGEKVPVIDYYCQTNNVPNHCGSPVL

>C.aristophanes_tpm:2463.91

MKTVAVFLVVAVAVAYGQFFCPDSEDDPLNCLETQATGATCMKSTEDDTYSYVCGYCGKKKASCFGKKVPSSDYNCQINKIPNHCGGPVM

>C.aristophanes_tpm:3200.72

MKTVAVFLVVAVAVAYGQFFCPDSEDDPLNCLETQATGATCMKSTEDDTYSYVCGYCGKKKASCFGKKVPTSDYNCQINKIPNHCGGPVM

>C.bayani_tpm:7477.39

MKTVAVFLVVALAVAYGQFFCPSSKDEPLNCIETMASTPTCMKSTADESLSYACGYCGKKKETCSGDKVPVSNYNCQIRKIPNPCGGPAL

>C.bayani_tpm:20.84

MKAVAVFLVVALAVAYGQFFCPESVNDPLNCLETMANSATCMQSNDDKSYSYVCGYCGKKKETCFGNKVAVLDYYCQTNGIANSCGGAAQ

>C.boavistensis_tpm:1307.04

MKTVAVFLVVALAVAYGQFFCPENESELNCIETMSTAATCMQSKDGSYSYACGYCGKKKDTCFGDKKPVRDYYCQRQKIVNNCGGPAL

>C.boavistensis_tpm:6.2

MKAVAVFMIVALAVAYGQFFCPESASDPLNCLETMASSATCMQSEADGSYSYVCGYCGKKKETCFGNKVAVRDYYCQTNGIANTCGGAAQ

>C.caracteristicus_tpm:9725.06

MKTVAVFLVVAIAVAYGQFFCPESISELSCVETMPNTATCMKSLDGSYSYACGYCGKKKEHCFGDKASVQDYYCQINSIANTCGGAAA

>C.consors_tpm:717.69

MKTVAVFLVVALAVAYGQFFCPDSEDDPLNCVETMATTATCMQSKGDKSFSYACGYCGKKKDSCFGDKVPVTDYHCKSRNIVNPCGGPAL

>C.consors_tpm:219.1

MKAVAVFMIVALAVAYGQFFCPESIDELNCLETQPNSAACMQSSDDDSYSYVCGYCGKKKEHCFGNKVAVQDYYCQTNGIANTCGGAA

>C.coronatus_tpm:1621.49

MKTVALFLVVAVAVAYGQFFCPDSEDDPLNCLETQATGATCMKSTEDDTYSYVCGYCGKKKASCFGKKVPSSDYNCQINKIPNHCGGPVM

>C.coronatus_tpm:15.67

MKAVAVFLVVALAVAYGQFFCPDSENDPLNCIETKATAATCMKSKTDGSYSYACGYCGKKKESCFGDKVPVTDYNCQTRKIANPCGGPAL

>C.cuneolus_tpm:5483.52

MKTVAVFLVVALAVAYGQFFCPENESELNCIETMSTAATCMQSKDGSYSYACGYCGKKKDTCFGDKKPVRDYYCQRQKIVNNCGGPAL

>C.ebraeus_tpm:1080.6

MKTVAVFLVVALAVAYGQFFCPDSEDDPLNCLETQATSATCMKSTEDETYSYVCGYCGKKKESCFGKKVPTSDYNCQIHKIPNHCGGPVM

>C.ebraeus_tpm:13017.92

MKTVAVFLVVALAVAYGQFFCPDSEDDPLNCLETQATSATCMKSTEDETYSYVCGYCGKKKEACFGKKVPSSDYNCQIHKIPNHCGGPVM

>C.ebraeus_tpm:43.86

MKAVAVFMIVALAVAYGQFFCPESENDPLNCLETQPNSATCMQSSDDQSYSYVCGYCGKKKETCFGNKAAVQDYYCQRNGIANTCGGAAQ

>C.ermineus_tpm:87030.59

MKTVAVFLVVALAVAYGQFFCPESKDEPLNCIETMASAATCMKSTTDGTFSYVCGYCGKKKESCSGEKVPVSDYSCQIHNVVNNCGGAAL

>C.ermineus_tpm:18.43

MKTVAVFLVVALAVAYGQFFCPENESELNCIETMSTAATCMQSKDGSYSYACGYCGKKKDTCFGDKKPVRDYYCQRQKIVNNCGGPAL

>C.galeao_tpm:1763.14

MKTVAVFLVVALAVAYGQFFCPENESDVNCIETMPTAVTCMQSKDGSYSYACGYCGKKKDTCFGDKKAVKDYYCQRQKIVNNCGGPAL

>C.galeao_tpm:9.49

MKAVAVFMIVALAVAYGQFFCPESASDPLNCLETMASSATCMQSEADGSYSYVCGYCGKKKETCFGNKVAVRDYYCQTNGIANTCGGAAQ

>C.generalis_tpm:9993.99

MKAVAVFLVVALAVAYGQFFCPKSKDDPLNCYETMPTKATCMKSKSGEIYTYACGYCGKKKESCAGENEPVADYFCRINKIPNPCGSPVL

>C.grahami_tpm:5991.35

MKTVAVFLVVALAVAYGQFFCPENESELNCIETMTTAATCMQSKDGSYSYACGYCGKKKDTCFGDKKPVRDYYCQRQKIVNNCGGPAL

>C.grahami_tpm:121.4

MKAVAVFMIVALAVAYGQFFCPESASDPLNCLETMASSATCMQSEADGSYSYVCGYCGKKKETCFGNKVAVRDYYCQTNGIANTCGGAAQ

>C.guanche_tpm:1488.96

MKTVAVFLVVALAVAYGQFFCPENENELNCIETMATAATCMQSKDGSYSYACGYCGKKKDTCFGDKTPVTDYYCQIQKIVNHCSGPAL

>C.guanche_tpm:12.16

MKVVAVFMIVALAVAYGQFFCPESASDSLNCLETMASSATCMQSNDDGSYSYVCGYCGKKKETCFGNKVGVQDYYCQTNGIANTCGGAAQ

>C.infinitus_tpm:1672.96

MKTVAVFLVVALAVAYGQFFCPENESELNCIETMSTAATCMQSKDGSYSYACGYCGKKKDTCFGDKKPVRDYYCQRQKIVNNCGGPAL

>C.infinitus_tpm:12.97

MKAVAVFMIVALAVAYGQFFCPESASDPLNCLETMASSATCMQSEADGSYSYVCGYCGKKKETCFGNKVAVRDYYCQTNGIANTCGGAAQ

>C.judaeus_tpm:165.87

MKTVAVFLVVALAVAYGQFFCPDSEDDPLNCLETQATAATCMKSTEDETYSYVCGYCGKKKETCFGKKVPTSDYNCQIHKIPNHCGGPVL

>C.lenavati_tpm:12443.79

MKAVAVFLVVALAVAYGQFFCPSSKDEPLNCIETMASTATCMKSTADESLSYACGYCGKKKETCSGDKVPVTNYNCQIKKIPNPCGGPAL

>C.lenavati_tpm:2610.79

MKAVAVFLVVALAVAYGQFFCPSSKDEPLNCIETMASTATCMKSTADESLSYACGYCGKKKETCSGDKVPVTNYNCQIKKIPNPCGGPAL

>C.lenavati_tpm:2297.02

MKAVAVFLVVALAVAYGQFFCPSSKDEPLNCIETMASTATCMKSTADESLSYACGYCGKKKETCSGDKVPVTNYNCQIKKIPNPCGGPAL

>C.litteratus_tpm:4905.91

MKVVAVFLVVALAVAYGQFFCPDSENDPLNCLETMASAATCMKSKSDGTYSYACGYCGKKKETCFGDKVPVTDYNCQRHKIVNHCGGPVV

>C.litteratus_tpm:13.95

MKAVAVFMIVALAVAYGQFFCPESVNDPLNCLESMPDTATCMRSTVDGSYSYACGYCGKKKEHCFGNKVGVKDYYCQRNGIVNTCGGAAQ

>C.lividus_tpm:4898.71

MKAVAVFLVVSLAVAYGQFFCPDSENDPLNCIETKASTATCMKSNTDHTYSYVCGYCGKKKETCFGNKVPVNDYHCQTNKIANNCGGAAL

>C.maioensis_tpm:20495.37

MKTVAVFLVVALAVAYGQFFCPENESELNCIETMSTAATCMQSKDGSYSYACGYCGKKKDTCFGDKKPVRDYYCQRQKIVNNCGGPAL

>C.maioensis_tpm:88.22

MKAVAVFMIVALAVAYGQFFCPESASDPLNCLETMASSATCMQSEADGSYSYVCGYCGKKKETCFGNKVAVRDYYCQTNGIANTCGGAAQ

>C.mercator_tpm:3099.08

MKTVAVFLVVALAMAYGQFFCPENENELNCIETMATAATCMQSKDGSYSYACGYCGKKKDTCFGDKTPVTDYYCQIQKIVNHCSGPAL

>C.miliaris_tpm:2344.53

MKTVAVFLLVAVAVAYGQFFCPDSEDDPLNCLETKGTQATCMKSTEDDTYSYVCGYCGQKKASCFGKKVPSSDYNCQINKIPNHCGGPVM

>C.miliaris_tpm:2147.99

MKTVAVFLVVALAVAYGQFFCPDSEDDPLNCLETKATGATCMKSTEDDTYSYVCGYCGQKKASCFGKKVPSSDYNCQIRKIPNHCGGPVM

>C.miruchae_tpm:21489.99

MKTVAVFLVVALAVAYGQFFCPENESELNCIETMSTAATCMQSKDGSYSYACGYCGKKKDTCFGDKKPVRDYYCQRQKIVNNCGGPAL

>C.miruchae_tpm:278.48

MKAVAVFMIVALAVAYGQFFCPESASDPLNCLETMASSATCMQSEADGSYSYVCGYCGKKKETCFGNKVAVRDYYCQTNGIANTCGGAAQ

>C.mordeirae_tpm:14615.65

MKTVAVFLVVALAVAYGQFFCPENENELNCIETMSTAATCMQSKDGSYSYACGYCGKKKDTCFGDKKPVRDYYCQRQKIVNNCGGPAL

>C.obscurus_tpm:16672.37

MKTVAVFLVVALAVAYGQFFCPSSKEESLNCIETMGTTATCMKTNKGDVYSYACGYCGKKKESCFGDKKPVTDYQCQTRNIPNPCGGAAR

>C.rattus_tpm:7476.74

MKAVAVFLVVAIAVAYGQFFCPGSKDDPLNCIETMGTTATCMKSTTDGSYSYACGYCGKKKESCFGNKVPVKDYYCQTSKVANNCGGTAL

>C.raulsilvai_tpm:3731.18

MKTVAVFLVVALAVAYGQFFCPENESELNCIETMSTAATCMQSKDGSYSYACGYCGKKKDTCFGDKKPVRDYYCQRQKIVNNCGGPAL

>C.raulsilvai_tpm:42.37

MKAVAVFMIVALAVAYGQFFCPESASDPLNCLETMASSATCMQSEADGSYSYVCGYCGKKKETCFGNKVAVRDYYCQTNGIANNCGGAAQ

>C.regonae_tpm:13011.28

MKTVAVFLVVALAVAYGQFFCPENESELNCIETMSTAATCMQSKDGSYSYACGYCGKKKDTCFGDKKPVRDYYCQRQKIVNNCGGPAL

>C.regonae_tpm:34.12

MKAVAVFMIVALAVAYGQFFCPESASDPLNCLETMASSATCMQSEADGSYSYVCGYCGKKKETCFGNKVAVRDYYCQTNGIANTCGGAAQ

>C.sponsalis_tpm:4229.31

MKTVAVLLLVSLAVAYGQFFCPDSENDPLNCIETMASSATCMKSKDGTHSYVCGYCGKKKESCFGGKVPVSDYNCQTRKIANPCGGAAL

>C.sponsalis_tpm:36.89

MKAVAVFMIVALAVAYGQFFCPESEADLNCLETQPNSAACMQSSDDGSYSYVCGYCGKKKEHCFGNKAAVQDYYCQTNGIANTCGGAA

>C.sulcunatus_tpm:13677.84

MKVVAVFLVVALAAAYGQFFCPDSENDPLNCVETMASAATCMKSRDGSYSYACGYCGKKKESCFGDKVPVTNYHCQTRKIPNKCGGPVL

>C.sulcunatus_tpm:96.48

MKAVAVFMIVALAVAYGQFFCPESVNDPLNCMETMPNTATCMQSSVDRSYSYACGYCGKKKETCFGNKAAVRDYFCQRNGIANPCGGAAQ

>C.terebra_tpm:3759.69

MKAVAVFLVVALAVAYGQFFCPDNENDPLNCVETKPFEATCMKSKDGTYSYVCGYCGTKNETCFGEKVPISDYSCKSRRVVNNCGGPVV

>C.tribblei_tpm:986.34

MKTVAVFLVVALAVAYGQFFCPSSKDEPLNCIETMASTATCMKSTTDESLSYACGYCGKKKETCSGDKVPVTNYNCQIKKIPNPCGGPAL

>C.trochulus_tpm:7535.08

MKTVAVFLVVALAVAYGQFFCPESEDDPLNCVETMATSATCMKSSDGSFSYACGYCGKKKETCFGNKQPVSDYYCQTQKIVNHCSGTVL

>C.trochulus_tpm:69.77

MKVVAVFMMVALAVAYGQFFCPESANDPLNCMETMASSATCMQSNADGSYSYVCGYCGKKKETCFGNKVAVQDYYCQTNSIANTCGGAAQ

>C.varius_tpm:5760.4

MKAVAVFLIVGLAVAYGQFFCPKNKDDPLNCVETMSTSASCFESEDGSYTYVCGYCGKKKASCSGEKKQVDDYYCRINKIANHCGSTAL

>C.verdensis_tpm:2723.9

MKTVAVFLVVALAVAYGQFFCPENESELNCIETMSTAATCMQSKDGSYSYACGYCGKKKDTCFGDKKPVRDYYCQRQKIVNNCGGPAL

>C.virgo_tpm:9721.95

MKAVAVFLVVAVAVAYGQFFCPDSENDPLNCLETMPSAATCMKSKSDGTYSYVCGYCGKKKETCFGDKVPVTDYSCQTRKVVNNCGGPVL
