## Supplementary material for "Conkazal-M1 from the MKAVA family of conotoxins – a dual-function protease inhibitor and neuroactive peptide": Suppl. File 3

```
Supplementary File 3. All sequences used for the CLANS clustering analysis in Fig. 5b.


>18400X15.C.geographus.TRINITY_DN533_c0_g1_i1#supFam_MKAVA#tpm_4696.52#len_270_dd0_1 0 0 0
MKTVAVFLVVALAVAYGQFFCPSSKEDSLNCIETMATTATCMKSNKGEIYSYACGYCGKKKESCFGDKKPVTDYQCQTRNIPNPCGGAAL
>SRR9831255.Pionoconus.magus.VG.TRINITY_DN6423_c0_g1_i1	supFam:MKAVA	tpm:12252	len:91 1 1 1
MKTVAVFLVVALAVAYGQFFCPNSENDPLNCVETMATTATCMQSNKDKSYSYACGYCGKKKESCFGNKVPVRDYNCKSRNIVNPCGGPAL
>13691X5.ImpB1.TRINITY_DN2865_c0_g1_i1_Entry:2525.conotoxin.MKAVA_len:498_tpm:6010.22 2 2 2
MKAVAVFLVVSLAVAYGQFFCPSSKNEPLNCLETMATTATCMKSTKDGTYSYVCGYCGKKKETCSGDKVPVTDYNCRVKNIANLCGGPLH
>14357X3.C.furvus.TRINITY_DN11037_c0_g1_i1_Entry:2525.conotoxin.MKAVA_len:548_tpm:5758.00 4 3 3
MKAVAVFLVVALAVAYGQFFCPDSEHDPLNCVETKAFAATCMKSKDGSYSYACGYCGKKKESCFGDKVPVTDYNCKTRKIPNPCGGPAL
>14357X4.C.cordigerus.TRINITY_DN2751_c0_g1_i1_Entry:2525.conotoxin.MKAVA_len:548_tpm:1349.43 5 4 4
MKAVAVFLVVALAVAYAQFFCPDNENDPLNCIETKASIATCMKSNTDGSFSYACGYCGKKKESCFGDKVPVTDYNCQTRNIANPCGGAAL
>14718X3.C.lenavati.TRINITY_DN5872_c0_g1_i1_Entry:2525.conotoxin.MKAVA_len:558_tpm:12443.79 6 5 5
MKAVAVFLVVALAVAYGQFFCPSSKDEPLNCIETMASTATCMKSTADESLSYACGYCGKKKETCSGDKVPVTNYNCQIKKIPNPCGGPAL
>C.imperialis.deep.VD.TRINITY_DN516_c0_g1_i1_Entry:2525.conotoxin.MKAVA_len:475_tpm:4787.72 10 6 6
MKAVAVFLVVSLAVAYGQFFCPSSKNEPLNCLETMASTATCMKSTKDGSYSYVCGYCGKKKETCSGDKVPVTDYNCRVNKIANLCGGAVL
>rolani.TRINITY_DN21766_c0_g1_i1_Entry:2525.conotoxin.MKAVA_len:573_tpm:4960.87 11 7 7
MKVVAVFLVVALAAAYGQFFCPDNENDPLNCIETMASGATCMKSNKDGSYSYACGYCGKKKESCFGDKVPVTNYHCQTRKIPNKCGGPVL
>SRR14407584.C.abbreviatus.TRINITY_DN301_c0_g1_i1_Entry:2525.conotoxin.MKAVA_len:491_tpm:1489.23 16 8 8
MKAVAVFLVVVIAVAYGQFFCPGSKDDPLNCIETMGTTATCMRSTTDGSYSYACGYCGKKKESCFGNKVPVKDYYCQTSKVANNCGGTAL
>SRR2124881.C.betulinus.VG.TRINITY_DN7727_c0_g1_i1_Entry:2525.conotoxin.MKAVA_len:494_tpm:5832.33 17 9 9
MKAVAVFLVVALAVAYGQFFCPESENDPLNCIETMASAATCMKSKADGSYSYACGYCGKKKESCFGDKVPVTDYNCKTRKIVNPCGGPAL
>gloriamaris.TRINITY_DN2395_c0_g2_i1_Entry:2525.conotoxin.MKAVA_len:544_tpm:2550.18 23 10 10
MKAVAVFLVVALAVAYGQFFCPDNENDPLNCVETQGSVATCMRSTLDKSHSYVCGYCGKKKESCFGNKVPVADYDCKMRKIANPCGSAR
>lividus.TRINITY_DN536_c0_g1_i1_Entry:2525.conotoxin.MKAVA_len:529_tpm:4898.71 24 11 11
MKAVAVFLVVSLAVAYGQFFCPDSENDPLNCIETKASTATCMKSNTDHTYSYVCGYCGKKKETCFGNKVPVNDYHCQTNKIANNCGGAAL
>quercinus.TRINITY_DN371_c0_g1_i1_Entry:2525.conotoxin.MKAVA_len:533_tpm:8305.29 26 12 12
MKAVAVFLVVALAVAYGQFFCPDSENDPLNCIETMASHATCMKSNTDGTYSYACGYCGKKKETCFGDKVPVTDYNCQTHKVVNNCGGAAL
>rattus.TRINITY_DN2275_c0_g1_i1_Entry:2525.conotoxin.MKAVA_len:521_tpm:7476.74 27 13 13
MKAVAVFLVVAIAVAYGQFFCPGSKDDPLNCIETMGTTATCMKSTTDGSYSYACGYCGKKKESCFGNKVPVKDYYCQTSKVANNCGGTAL
>terebra.TRINITY_DN1109_c0_g2_i1_Entry:2525.conotoxin.MKAVA_len:508_tpm:1884.26 31 14 14
MKAVAVFLVVALAVAYGQFFCPDNENDPLNCVETKPFEATCMKSKDGTYSYVCGYCGTKNETCFGEKVPISDYSCKSRRVVNNCGGPVV
>textile.TRINITY_DN5626_c0_g1_i1_Entry:5671.conotoxin.MKAVA_len:545_tpm:6208.73 32 15 15
MKAVAVFLVVALAVAYGQFFCPDSENDPLNCVETMGTLATCMKSKSGKSYSYACGYCGKKKESCFGDKMPVDAYDCQVRNIANPCGGSAL
>varius_MP.TRINITY_DN17_c0_g1_i1_Entry:2525.conotoxin.MKAVA_len:507_tpm:5760.40 34 16 16
MKAVAVFLIVGLAVAYGQFFCPKNKDDPLNCVETMSTSASCFESEDGSYTYVCGYCGKKKASCSGEKKQVDDYYCRINKIANHCGSTAL
>virgo_MP.TRINITY_DN558_c0_g1_i1_Entry:2525.conotoxin.MKAVA_len:540_tpm:9721.95 35 17 17
MKAVAVFLVVAVAVAYGQFFCPDSENDPLNCLETMPSAATCMKSKSDGTYSYVCGYCGKKKETCFGDKVPVTDYSCQTRKVVNNCGGPVL
>NP_001191584.1 schistosomin-like precursor [Aplysia californica] 88 18 18
MMRQLFVLVIVAGAVLADNYWCPKAGEAFECFESSPTERFCLTNGRETAVICSKCRKKFDFCRNDFVKSKRAHTDCGAGWESTPCTHDNSHVPAVFPGKL
>XP_005098364.1 schistosomin isoform X1 [Aplysia californica] 89 19 19
MMRQLFVLVIVAGAVLADNYWCPKAGEAFECFESSPTERFCLTNGRETAVICSKCKKKFDFCRNDFVKSKRADTDCGAGWESTPCTHDNSHVPAVFPGKL
>RUS82021.1 hypothetical protein EGW08_010212 [Elysia chlorotica] 90 20 20
MAPQVTNRSDHRPIYLGPLDMDWSHQPKRSRHRRSKSLHTSHCSNNCHYNSAKMMKQLVIIAALACAVLADNYNCGKKGDVFECFESAAADRFCLTGGAAPVVVCGKCRKKADYCTKGLKKSSRPEVDCGPDFASTPCTTANSAVPAFF
>KAH9500249.1 hypothetical protein Btru_077571 [Bulinus truncatus] 91 21 21
MKTVFLILALIVCAAMADNYRCVNPGEPFECFESDATARFCVSGNKGAYVICSKCRRKYEFCANGAKVSKRPEVECGANWASTDCTSDNSDVPSVL
>NP_001298209.1 schistosomin-like precursor [Biomphalaria glabrata] 92 22 22
MKTVFFILALIVCAVVADNYRCPNPGDAFECFESDATARFCVSGKRGAYVICSKCRRKYEFCANGAKVSKRPEVECRADWASTECTSDNSDVPSVM
>GFO13819.1 schistosomin-like [Plakobranchus ocellatus] 94 23 23
MKQIVIILAVACVAVLADNYMCGKRGAKFECFESAASDRFCLTGGAHPEVVCGKCRKKADFCTKGLVMSKRPAIDCGASYESTPCTTGNSEVPATY
>GFS27466.1 schistosomin-like [Elysia marginata] 95 24 24
MAPKLWQLSFKTNSYDEQNMQRELTFLSFLSFSPLMSGFTAETTQYSHGCEFNWLLWCAYRDPDCRCSSIFTDSQYRYNLASVKMMKQIVLLAAIACAAVLADNYNCGNKGDKFECFESAASARFCLTGGAQPEVICGKCRKKADFCTKGLVLSKRPEVDCGAGYASTPCTTANSAVPATF
>sp|P24471|SCHS_LYMST Schistosomin OS=Lymnaea stagnalis OX=6523 PE=1 SV=1 96 25 25
DNYWCPQSGEAFECFESDPNAKFCLNSGKTSVVICSKCRKKYEFCRNGLKVSKRPDYDCGAGWESTPCTGDNSAVPAVF
>tr|A0A2T7P5S2|A0A2T7P5S2_POMCA Uncharacterized protein OS=Pomacea canaliculata OX=400727 GN=C0Q70_11377 PE=4 SV=1 98 27 26
MKIIAALALLAVAVHARFSCPPDPTQGIDCFETEPDQLFCMSDGNLVCGTCGMRLAFCLGLVEGVIIIQTNDPAVCAGAIPPECPTGNSQQRPGGQGGQGGHGGHGNPFGIPPGLNKPKN
>GICZ01142588.1 TSA: Berthella plumula breed wildtype TRINITY_DN16772_c0_g1_i2, transcribed RNA sequence 99 28 27
MMKEIVFLALLACVMADNYLCPNDGDRFDCFESAPNDRFCLTDGRVHKVVCSKCRKKYDFCETEGDFHESKTPEKHCGDDYEATPCTRQNSHVPAKLKIIKRPL
>GGMZ01015001.1 TSA: Phylliroe bucephala Phylliroe_bucephala_comp12891_c0_seq1 transcribed RNA sequence 100 29 28
MMKIFILLVVVCTAVMGDNYLCPKEGVKYECEEYRADKPVCLSNGKESKVVCGFCRQKLEFCRSGFEKSKHSERHCGEGWENTPCTPKNSAVPEFISYN
>GIYH01067931.1 TSA: Berghia stephanieae TRINITY_DN281388_c0_g1_i1, transcribed RNA sequence 101 30 29
MKTILCLLLIGCAAVLADKYLCPVKDGELFECFESDPEYRGCLTNGRRSESYVICTKCKRKWEFCVTRGAFSVSKHPDTDCGADYKNTPCTPQNSEWSSVFKFRV
>GIBV01002871.1 TSA: Eubranchus exiguus breed wildtype BINPACKER.1025.6, transcribed RNA sequence 102 31 30
MNSTAILSFIALFAFATVLADNYLCPSADGEAFECFESQADHRTCLTNKRRTATYVVCGKCTKKWEFCVTKGEMLTSKRPERDCGADYESTPCTAANSFWDKSFKYRSN
>GICX01048853.1 TSA: Cratena peregrina breed wildtype Single_4233, transcribed RNA sequence 103 32 31
MKIVLCLLFIGVAIVLADNYLCPTRDGEPFECLESDVTYRSCLTNGRRGDAYVVCSKCKMKWEFCVTRGTFSESKHPESDCGADYKTTPCTTENSEWAKTFKLRV
>GIDA01088326.1 TSA: Hancockia uncinata breed wildtype Shannon_HW16_H_cfuncinata_c1_57_9270_927, transcribed RNA sequence 104 33 32
MKATLFLLLIGCGTILCAKFICPKEGEDFECFEFNPEYRACMTDGRKANSYVTCSKCKVKWEYCRTGGKFEEEKKHADRYCPPDYKSTPCTPENSHVGKVFTKA
>GJHI01073086.1 TSA: Atlanta ariejansseni TRINITY_DN31_c1_g1_i16, transcribed RNA sequence 105 34 33
MKTVAILLLVIAAASAQFFCPKKGDPIQCYETRPHTAFCMANGDWVCGKCTVKTLSCQSGLRKGTRCSPSEKPAC
>GAQQ01020425.1 TSA: Bithynia siamensis goniomphalos CL8781.Contig1_All transcribed RNA sequence 106 35 34
MKTFVVLAMLVAVAYCQYFCPGPGEALECLDGPQSAAKAFCLSDGTWVCGNCKAKEIYCSSGVLKARFESTCAGNPKPDCKSQ
>GJGI01024251.1 TSA: Alviniconcha strummeri TRINITY_DN24261_c0_g1_i1, transcribed RNA sequence 107 36 35
MKTIAILALLVAAACAQYFCPDSEEQFQCLETKPNTAFCMTDTTWVCGKCAVKRSVCFGSATQAKHETVCTNAGNPKITC
>GGFE01018530.1 TSA: Potamopyrgus antipodarum cds.comp77318_c0_seq1.m.18001 transcribed RNA sequence 108 37 36
MKTFIVLAMLVAVAYSQFFCNDDGSAMECMDGPVTAATAYCLNNGNWVCGKCAVKEIFCSSGAAIKQSHTEQTCTNAGNPKPSC
>GDIA01100366.1 TSA: Rapana venosa c117256_g1_i1 transcribed RNA sequence 109 38 37
MKVAILFLAIAVVAVYGQYFFCPPGESAEECSFNYNLSNPACMKDLDGSGTLSYICTYCEVKAASCSGEKEQVDSSNCQNSNVNNPCP
>GCEL01081520.1 TSA: Cipangopaludina cathayensis CCAH01081899 transcribed RNA sequence 110 39 38
MKCFVVLALLVAVVGAIFFCPDSEDQLQCLETQPATAYCLTNGQWVCGKCGIKKQNCFGLTDRQGRRPVEDRSKNACYGKRVPSCGQ
>GHAT01224603.1 TSA: Crassostrea gigas TR82664_c0_g1_i1, transcribed RNA sequence 111 40 39
MRKELIILAVVVCAVLADNYWCPQSGEAFECFESDPNAKFCLNSGKTSVVICSKCRKKYEFCRNGLKVSKRPDYDCGAGWESTPCTGDNSAVPAVF
>XP_025098673.1 uncharacterized protein LOC112566623 [Pomacea canaliculata] 114 43 40
MKIIVALALLAVAVLAYECPPDPTQGIDCFETEPGKVFCMTDGSLVCGVCGKRLAYCLGMMEGLLILESIDPSVCEGATAPECEQGISGPKIKG
>PVD28780.1 hypothetical protein C0Q70_11375 [Pomacea canaliculata] 115 44 41
MKCIVLLALLVVAANTRFICPSDLSTFQCFETAPGVPMCLTDYTWVCGNCGVKWAYCEAKADGVNNPPIKTSDAVCAGTPKPQC
>18400X15.C.geographus.TRINITY_DN533_c0_g1_i1#supFam_MKAVA#tpm_4696.52#len_270_dd0_1 0 49 42
MKTVAVFLVVALAVAYGQFFCPSSKEDSLNCIETMATTATCMKSNKGEIYSYACGYCGKKKESCFGDKKPVTDYQCQTRNIPNPCGGAAL
>SRR9831255.Pionoconus.magus.VG.TRINITY_DN6423_c0_g1_i1	supFam:MKAVA	tpm:12252	len:91 1 50 43
MKTVAVFLVVALAVAYGQFFCPNSENDPLNCVETMATTATCMQSNKDKSYSYACGYCGKKKESCFGNKVPVRDYNCKSRNIVNPCGGPAL
>13691X5.ImpB1.TRINITY_DN2865_c0_g1_i1_Entry:2525.conotoxin.MKAVA_len:498_tpm:6010.22 2 51 44
MKAVAVFLVVSLAVAYGQFFCPSSKNEPLNCLETMATTATCMKSTKDGTYSYVCGYCGKKKETCSGDKVPVTDYNCRVKNIANLCGGPLH
>13691X5.ImpB1.TRINITY_DN1537_c0_g1_i1_Entry:2525.conotoxin.MKAVA_len:470_tpm:61.93 3 52 45
MKAVAVFLVVSLAVAYGQFFCPASVNDPLNCLETMPNSATCMQFSGDGSYSYVCGYCGKKNETCFGNKVAVNDYYCQTKNIANNCGGAAQ
>14357X3.C.furvus.TRINITY_DN11037_c0_g1_i1_Entry:2525.conotoxin.MKAVA_len:548_tpm:5758.00 4 53 46
MKAVAVFLVVALAVAYGQFFCPDSEHDPLNCVETKAFAATCMKSKDGSYSYACGYCGKKKESCFGDKVPVTDYNCKTRKIPNPCGGPAL
>14357X4.C.cordigerus.TRINITY_DN2751_c0_g1_i1_Entry:2525.conotoxin.MKAVA_len:548_tpm:1349.43 5 54 47
MKAVAVFLVVALAVAYAQFFCPDNENDPLNCIETKASIATCMKSNTDGSFSYACGYCGKKKESCFGDKVPVTDYNCQTRNIANPCGGAAL
>14718X3.C.lenavati.TRINITY_DN5872_c0_g1_i1_Entry:2525.conotoxin.MKAVA_len:558_tpm:12443.79 6 55 48
MKAVAVFLVVALAVAYGQFFCPSSKDEPLNCIETMASTATCMKSTADESLSYACGYCGKKKETCSGDKVPVTNYNCQIKKIPNPCGGPAL
>14718X3.C.lenavati.TRINITY_DN16852_c0_g1_i1_Entry:2525.conotoxin.MKAVA_len:389_tpm:13.78 7 56 49
MKAVAVFLVVALAVAYGQFFCPESVNDPLNCLETMPNSATCMQSSVDNSYSYVCGYCGKKKETCFGNKVAVQEYYCQTNGIANSCGGGAQ
>14718X8.C.litteratus.TRINITY_DN7827_c0_g1_i1_Entry:2525.conotoxin.MKAVA_len:466_tpm:13.95 8 57 50
MKAVAVFMIVALAVAYGQFFCPESVNDPLNCLESMPDTATCMRSTVDGSYSYACGYCGKKKEHCFGNKVGVKDYYCQRNGIVNTCGGAAQ
>18400X15.C.geographus.TRINITY_DN6629_c0_g1_i1_Entry:2525.conotoxin.MKAVA_len:483_tpm:22.59 9 58 51
MKAVAVFMIVALAVAYGQFFCPESVNDPLNCLESQPNSATCMQSSLDNSYSYVCGYCGKKKETCFGNKVAVQDYYCQTNGIANTCGGAA
>C.imperialis.deep.VD.TRINITY_DN516_c0_g1_i1_Entry:2525.conotoxin.MKAVA_len:475_tpm:4787.72 10 59 52
MKAVAVFLVVSLAVAYGQFFCPSSKNEPLNCLETMASTATCMKSTKDGSYSYVCGYCGKKKETCSGDKVPVTDYNCRVNKIANLCGGAVL
>rolani.TRINITY_DN21766_c0_g1_i1_Entry:2525.conotoxin.MKAVA_len:573_tpm:4960.87 11 60 53
MKVVAVFLVVALAAAYGQFFCPDNENDPLNCIETMASGATCMKSNKDGSYSYACGYCGKKKESCFGDKVPVTNYHCQTRKIPNKCGGPVL
>C.imperialis.shallow.VD.TRINITY_DN735_c0_g1_i1_Entry:2525.conotoxin.MKAVA_len:461_tpm:6.18 12 61 54
MKAVAVFLVVSLAVAYGQFFCPASVNDPLNCLETMPNSATCMQSNGDGTYSYVCGYCGKKNETCFGNKVAVNDYYCQTNSIANNCGGAAQ
>SRR11807492.C.raulsilvai.VG.TRINITY_DN5501_c0_g1_i1_Entry:2525.conotoxin.MKAVA_len:466_tpm:21.19 13 62 55
MKAVAVFMIVALAVAYGQFFCPESASDPLNCLETMASSATCMQSEADGSYSYVCGYCGKKKETCFGNKVAVRDYYCQTNGIANNCGGAAQ
>SRR13740844.C.ventricosus.TRINITY_DN14976_c0_g1_i1_Entry:2525.conotoxin.MKAVA_len:469_tpm:9.61 14 63 56
MKAVAVFMIVALAVAYGQFFCPESASAPLNCLETMASSATCMQSEVDGSYSYVCGYCGKKKETCFGNKKAVRDYYCQTNGIANTCGGAAQ
>SRR13781584.C.bayani.VG.TRINITY_DN1262_c0_g1_i1_Entry:2525.conotoxin.MKAVA_len:480_tpm:20.84 15 64 57
MKAVAVFLVVALAVAYGQFFCPESVNDPLNCLETMANSATCMQSNDDKSYSYVCGYCGKKKETCFGNKVAVLDYYCQTNGIANSCGGAAQ
>SRR14407584.C.abbreviatus.TRINITY_DN301_c0_g1_i1_Entry:2525.conotoxin.MKAVA_len:491_tpm:1489.23 16 65 58
MKAVAVFLVVVIAVAYGQFFCPGSKDDPLNCIETMGTTATCMRSTTDGSYSYACGYCGKKKESCFGNKVPVKDYYCQTSKVANNCGGTAL
>SRR2124881.C.betulinus.VG.TRINITY_DN7727_c0_g1_i1_Entry:2525.conotoxin.MKAVA_len:494_tpm:5832.33 17 66 59
MKAVAVFLVVALAVAYGQFFCPESENDPLNCIETMASAATCMKSKADGSYSYACGYCGKKKESCFGDKVPVTDYNCKTRKIVNPCGGPAL
>SRR2124881.C.betulinus.VG.TRINITY_DN5828_c0_g1_i1_Entry:2525.conotoxin.MKAVA_len:373_tpm:168.13 18 67 60
MKAVAVFMIVALAVAYGQFFCPESVNDPLNCLETMASSATCMQSSVDGSYSYVCGYCGKKKETCFGNKVAVQDYYCQTNGIANSCGGAA
>SRR9831255.Pionoconus.magus.VG.TRINITY_DN4320_c0_g1_i1_Entry:2525.conotoxin.MKAVA_len:497_tpm:37.45 19 68 61
MKAVAVFMIVALAVAYGQFFCPESIDDLNCLETQPNSVACMQSSDDDSYSYVCGYCGKKKEHCFGNKVAVQDYYCQTNGIANTCGGAA
>consors.TRINITY_DN4789_c0_g1_i1_Entry:2525.conotoxin.MKAVA_len:512_tpm:219.10 20 69 62
MKAVAVFMIVALAVAYGQFFCPESIDELNCLETQPNSAACMQSSDDDSYSYVCGYCGKKKEHCFGNKVAVQDYYCQTNGIANTCGGAA
>coronatus.TRINITY_DN123_c0_g1_i1_Entry:2525.conotoxin.MKAVA_len:442_tpm:12.70 21 70 63
MKAVAVFLVVALAVAYGQFFCPDSENDPLNCIETKATAATCMKSKTDGSYSYACGYCGKKKESCFGDKVPVTDYNCQTRKIANPCGGPAL
>ebraeus.TRINITY_DN249_c0_g1_i1_Entry:2525.conotoxin.MKAVA_len:351_tpm:43.86 22 71 64
MKAVAVFMIVALAVAYGQFFCPESENDPLNCLETQPNSATCMQSSDDQSYSYVCGYCGKKKETCFGNKAAVQDYYCQRNGIANTCGGAAQ
>gloriamaris.TRINITY_DN2395_c0_g2_i1_Entry:2525.conotoxin.MKAVA_len:544_tpm:2550.18 23 72 65
MKAVAVFLVVALAVAYGQFFCPDNENDPLNCVETQGSVATCMRSTLDKSHSYVCGYCGKKKESCFGNKVPVADYDCKMRKIANPCGSAR
>lividus.TRINITY_DN536_c0_g1_i1_Entry:2525.conotoxin.MKAVA_len:529_tpm:4898.71 24 73 66
MKAVAVFLVVSLAVAYGQFFCPDSENDPLNCIETKASTATCMKSNTDHTYSYVCGYCGKKKETCFGNKVPVNDYHCQTNKIANNCGGAAL
>magus.TRINITY_DN6218_c0_g1_i1_Entry:2525.conotoxin.MKAVA_len:509_tpm:666.92 25 74 67
MKAVAVFMIVALAVAYGQFFCPESIDELNCLETQPNSVACMQSSDDNSYSYVCGYCGKKKEHCFGNKVAVQDYYCQTNGIANTCGGAA
>quercinus.TRINITY_DN371_c0_g1_i1_Entry:2525.conotoxin.MKAVA_len:533_tpm:8305.29 26 75 68
MKAVAVFLVVALAVAYGQFFCPDSENDPLNCIETMASHATCMKSNTDGTYSYACGYCGKKKETCFGDKVPVTDYNCQTHKVVNNCGGAAL
>rattus.TRINITY_DN2275_c0_g1_i1_Entry:2525.conotoxin.MKAVA_len:521_tpm:7476.74 27 76 69
MKAVAVFLVVAIAVAYGQFFCPGSKDDPLNCIETMGTTATCMKSTTDGSYSYACGYCGKKKESCFGNKVPVKDYYCQTSKVANNCGGTAL
>rolani.TRINITY_DN10411_c0_g1_i1_Entry:5671.conotoxin.MKAVA_len:473_tpm:26.18 28 77 70
MKAVAVFMIVALAVAYGQFFCPESVNDPLNCMETMPNTATCMKSSVDESYSYACGYCGKKKETCFGNKAAVRDYFCQRNGIANPCGGAAQ
>sponsalis.TRINITY_DN2091_c0_g1_i1_Entry:2525.conotoxin.MKAVA_len:468_tpm:36.89 29 78 71
MKAVAVFMIVALAVAYGQFFCPESEADLNCLETQPNSAACMQSSDDGSYSYVCGYCGKKKEHCFGNKAAVQDYYCQTNGIANTCGGAA
>striatus.TRINITY_DN13690_c0_g1_i1_Entry:2525.conotoxin.MKAVA_len:504_tpm:610.27 30 79 72
MKAVAVFMIVALAVAYGQFFCPESINELNCVETQPNSAACMQSSDDNSYSYVCGYCGKKKEHCFGNKVGVQDYYCQTNGIANTCGGAA
>terebra.TRINITY_DN1109_c0_g2_i1_Entry:2525.conotoxin.MKAVA_len:508_tpm:1884.26 31 80 73
MKAVAVFLVVALAVAYGQFFCPDNENDPLNCVETKPFEATCMKSKDGTYSYVCGYCGTKNETCFGEKVPISDYSCKSRRVVNNCGGPVV
>textile.TRINITY_DN5626_c0_g1_i1_Entry:5671.conotoxin.MKAVA_len:545_tpm:6208.73 32 81 74
MKAVAVFLVVALAVAYGQFFCPDSENDPLNCVETMGTLATCMKSKSGKSYSYACGYCGKKKESCFGDKMPVDAYDCQVRNIANPCGGSAL
>tribblei.all.TRINITY_DN3798_c0_g2_i1_Entry:2525.conotoxin.MKAVA_len:398_tpm:3.02 33 82 75
MKAVAVFLVVALAVAYGQFFCPESVNDPLNCLETMPNSATCMQSSDDKSYSYVCGYCGKKKETCFGNKVAVLDYYCQTNGIANSCGGGAQ
>varius_MP.TRINITY_DN17_c0_g1_i1_Entry:2525.conotoxin.MKAVA_len:507_tpm:5760.40 34 83 76
MKAVAVFLIVGLAVAYGQFFCPKNKDDPLNCVETMSTSASCFESEDGSYTYVCGYCGKKKASCSGEKKQVDDYYCRINKIANHCGSTAL
>virgo_MP.TRINITY_DN558_c0_g1_i1_Entry:2525.conotoxin.MKAVA_len:540_tpm:9721.95 35 84 77
MKAVAVFLVVAVAVAYGQFFCPDSENDPLNCLETMPSAATCMKSKSDGTYSYVCGYCGKKKETCFGDKVPVTDYSCQTRKVVNNCGGPVL
```
